# Pair-Living Emerged Early in Placental Mammals

**DOI:** 10.1101/2025.07.27.667013

**Authors:** S.F. Walmsley, C.A. Olivier, A. V. Jaeggi, L. Makuya, J. S. Martin, L.D. Hayes, C. Schradin

## Abstract

It is widely assumed that the first placental mammals were solitary. This assumption has been used as the default in previous comparative studies, but has never been tested independently. Here, we compiled a comprehensive database by reviewing over 14,000 primary peer-reviewed publications on the social organization of the 5,386 extant placental mammal species. We found empirical data for 738 species across 1,478 populations: 354 populations were primarily solitary, 241 populations pair-living and 878 populations group-living. Notably, 383 species (52%) and 580 populations (39%) exhibited intra-specific variation in social organization. Using a Bayesian phylogenetic framework that incorporates intra-specific variation, we show that the ancestral placental mammal - living approximately 160 to 100 million years ago - was not exclusively solitary, but likely displayed intraspecific variation. Pair-living was predicted to account for approximately 26% of ancestral social organisation. Our analysis found that ecology and life history of extant solitary and pair-living mammals was very similar and different from those of group-living mammals. Our results revise prevailing theories of mammalian social evolution, revealing that pair-living emerged early, shifting the focus of mammalian social evolution from the origins of pair-living to the origins of group-living.

## Main Text

Mammalian species display a high variety of social systems, ranging from solitary living to large groups with multiple adult males and females ^1^. It has long been assumed that this social diversity evolved from a solitary ancestor ^2–5^. However, recent comparative studies have found that pair-living was ancestral across different mammalian taxa^6,7^, and present in combination with solitary living ^8,9^. The solitary ancestry of mammals has not been tested in previous comparative studies ^2^. Thus, our understanding of the ancestral social organisation in early mammals remains incomplete, particularly regarding whether it was predominantly solitary and to what degree it was variable.

Previous studies on mammalian social evolution focused on understanding the evolution of pair-living ^2–5^. However, these studies might have asked the wrong question as the ancestors of elephant-shrews ^6^, artiodactyla ^7^ and primates ^8^ were already pair-living. Indeed, solitary living was often assumed as the *a priori* starting point of mammalian social evolution ^2,4^. Compounding this problem, hundreds of cryptic, nocturnal small species that have never been studied in the field have been assumed to be solitary in comparative databases ^2^, leading to biased estimates ^10^. To date, no comparative study has directly tested whether the ancestral mammal was solitary. Given fossil evidence that ancestors of mammals and early mammals— including metatherians and therians—lived in pairs or groups ^11–14^, this assumption should not be made *a priori* but rather tested *a posteriori*.

Studies on mammalian social evolution often focused on the origins of social monogamy ^2,4^. The concept of social monogamy combines the four components of social systems ^15^: social organisation (pair-living), social structure (pair bond), mating system (monogamy), and care system (biparental care). Each of these components varies considerably between pair-living species ^16^. It is, therefore, now widely accepted that analysing one single component of social systems yields clearer insights ^17–19^. Most available data concern social organisation, defined as the composition of social units, such as solitary, pair, or groups ^15^. Furthermore, while previous comparative studies assigned a single form of social organisation to each species ^2,4^, we know now that intra-specific variation in social organisation is common in mammals ^20^. This variation must be considered to accurately estimate ancestral social systems ^5,8^.

Various ecological and life-history factors have been proposed to drive transitions to social systems ^2,4,5,21^, and are central to socio-ecological models predicting social organization ^22–24^. Open habitats increase visibility to predators, selecting for group-living and larger body size^25–27^. Habitats with abundant, shareable, and defensible resources promote sociality^28,29^. Seasonal food scarcity affects breeding and social variation ^20,30^. Socio-ecological models have been criticized for overlooking life-history traits ^4,31^. Traits such as body size, diet, activity pattern, and sexual dimorphism correlate with sociality, but vary by taxon. For instance, larger primates tend to be social ^8^, but most large carnivores are solitary^32^. Sexual dimorphism is often greatest in group-living species due to male–male competition, but exceptions exist in solitary carnivores ^33^. Nocturnal species are often solitary, while diurnal species may form groups due to greater predation risk ^8^. Species on the slow end of the life-history continuum—long-lived, low-fecundity mammals—are more likely to be social ^34^. Together, ecological and life-history factors help explain interspecific variation in mammalian sociality.

Here we present the first study on the evolution of social organization in placental mammals that incorporates intra-specific variation and relies solely on peer reviewed data from field studies, excluding unverified assumptions about unstudied species. First, we investigated the extent to which different environmental and life history factors (Table S1) are associated to the different forms of social organisation that can be found in mammals (Table S2). Then, considering the increasing evidence for a socially flexible pair-living ancestor in many mammalian orders, we estimated the social organisation of the ancestor of all placentals and in how far it showed intra-specific variation in social organisation, expecting that it was socially flexibly and more sociable than previously assumed.

## Results

### Distribution of Social Organization in Extant Placentals

We searched primary literature for empirical data on the social organization of free-living placental mammals, reviewing approximately 14,000 articles. We extracted useable data from 2048 publications, covering 1478 populations from 738 of the 5386 extant placental species (Fig. 1; Table S3; Figure S1). The most common social organization of placental mammals (Figure 2A) was multi-male multi-female groups (MMFF; 483 populations, 32.8%) followed by solitary living (354 populations, 24.0%), one-male multi-female groups (MFF; 271 populations, 18.4%) and pair-living (MF; 241 populations = 16.4%). Sex-specific groups (84 populations = 5.7%), one-female multi-male groups (MMF; 18 populations = 1.2%) and sex-specific solitary male (22 populations = 1.49%) were rare (Figure 2B). Among studies that reported at least two social units, more than half (54.7%) showed intra-specific variation in social organization (IVSO), though there was substantial variation across populations (Fig. 2B).

**Figure 1.**
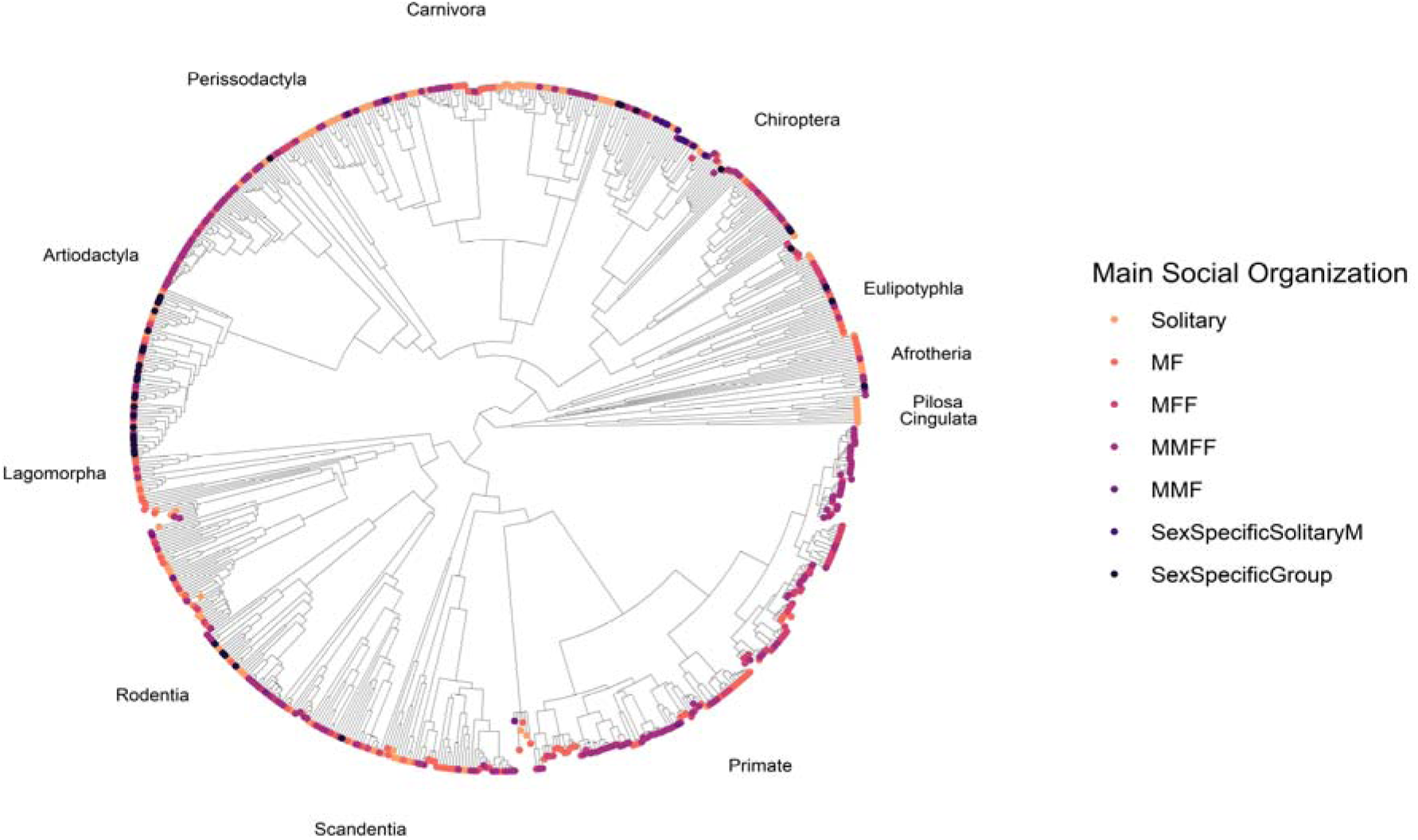
Phylogenetic tree of placental mammals included in this study. Orders of the Afrotheria superorder have been combined under a single label for visualization purposes (Proboscidea, Hyracoidea, Afrosoricida, Macroscelidea, Sirenia, Tubulidentata). Here we show the most common social organization summarized for each species. Solitary: both males and females are solitary; MF: pair-living (one male and one female); MFF: one male multi-female group; MMFF: multi-male multi-female groups; MMF: multi-males one female group; SexSM: Sex-specific with females in groups and males being solitary; SexSG: Sex-specific group: females in groups and males in other groups.

**Figure 2.**
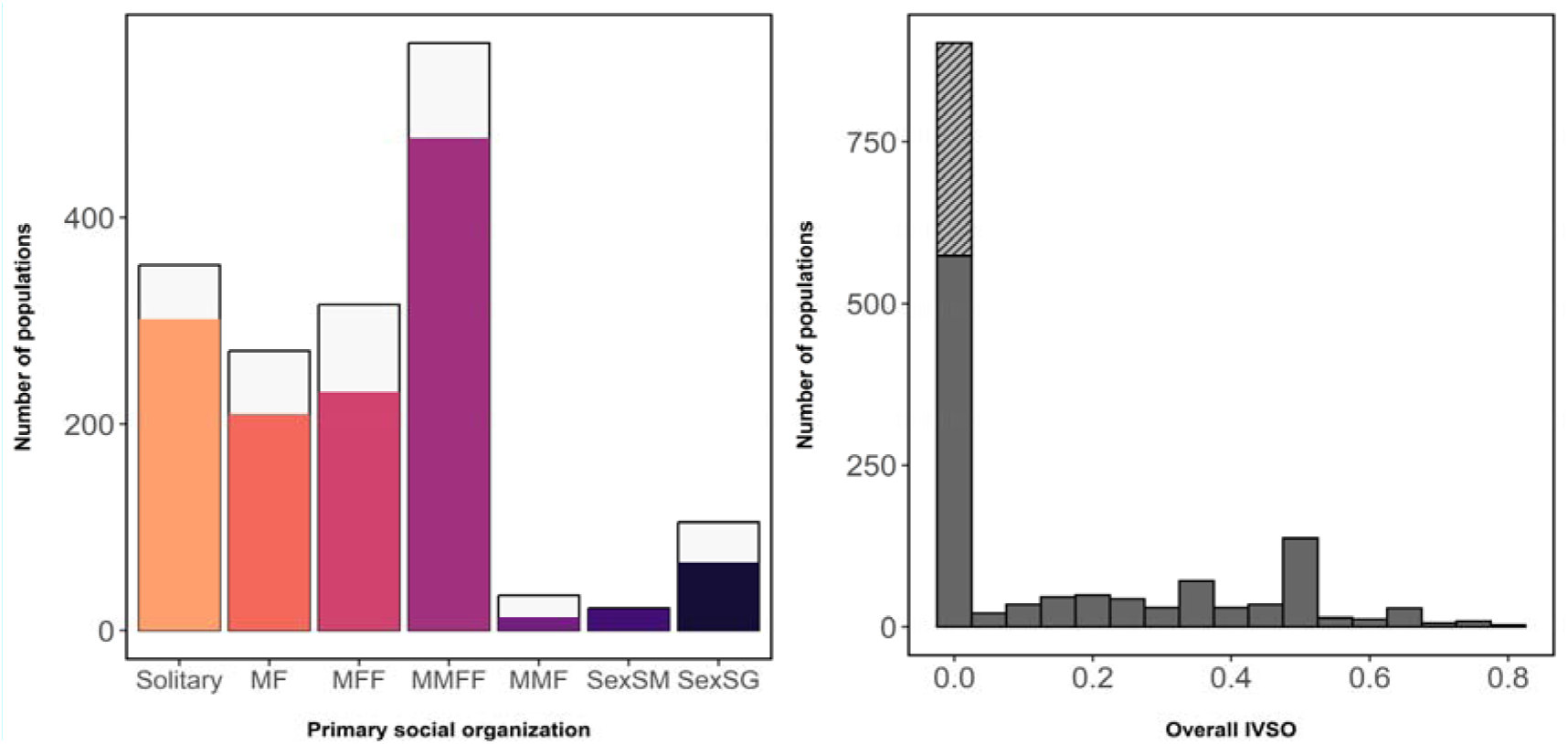
Left: Total number of populations in our dataset exhibiting each form of social organization as the primary (most common form). Light parts of the bars represent uncertainty in the main social organization for populations exhibiting two or more forms of social organization with equally high frequency (for example 50% of social units are solitary, 50% are pair-living). Solitary: both males and females are solitary; MF: pair-living (one male and one female); MFF: one male multi-female group; MMFF: multi-male multi-female groups; MMF: multi-males one female group; SexSM: Sex-specific with females in groups and males being solitary; SexSG: Sex-specific group: females in groups and males in other groups. Right: Total number of populations in our dataset exhibiting intra-specific variation in social organization (IVSO). Gray hashed part indicates populations where IVSO could not possibly be observed because only one group (social unit) was studied.

### Effects of ecological and life history predictors on social organization

The availability of measures for each predictor varied, such that sample size ranged from 848-1439 populations across 356-705 species (Table S4). Overall, the models showed strong associations between ecological or life history measures and social organization (Figures 3 & 4). Instead of emphasizing differences from the reference category (i.e., regression slopes), we highlight how absolute probabilities of each social organisation type changed with predictor values.

**Figure 3.**
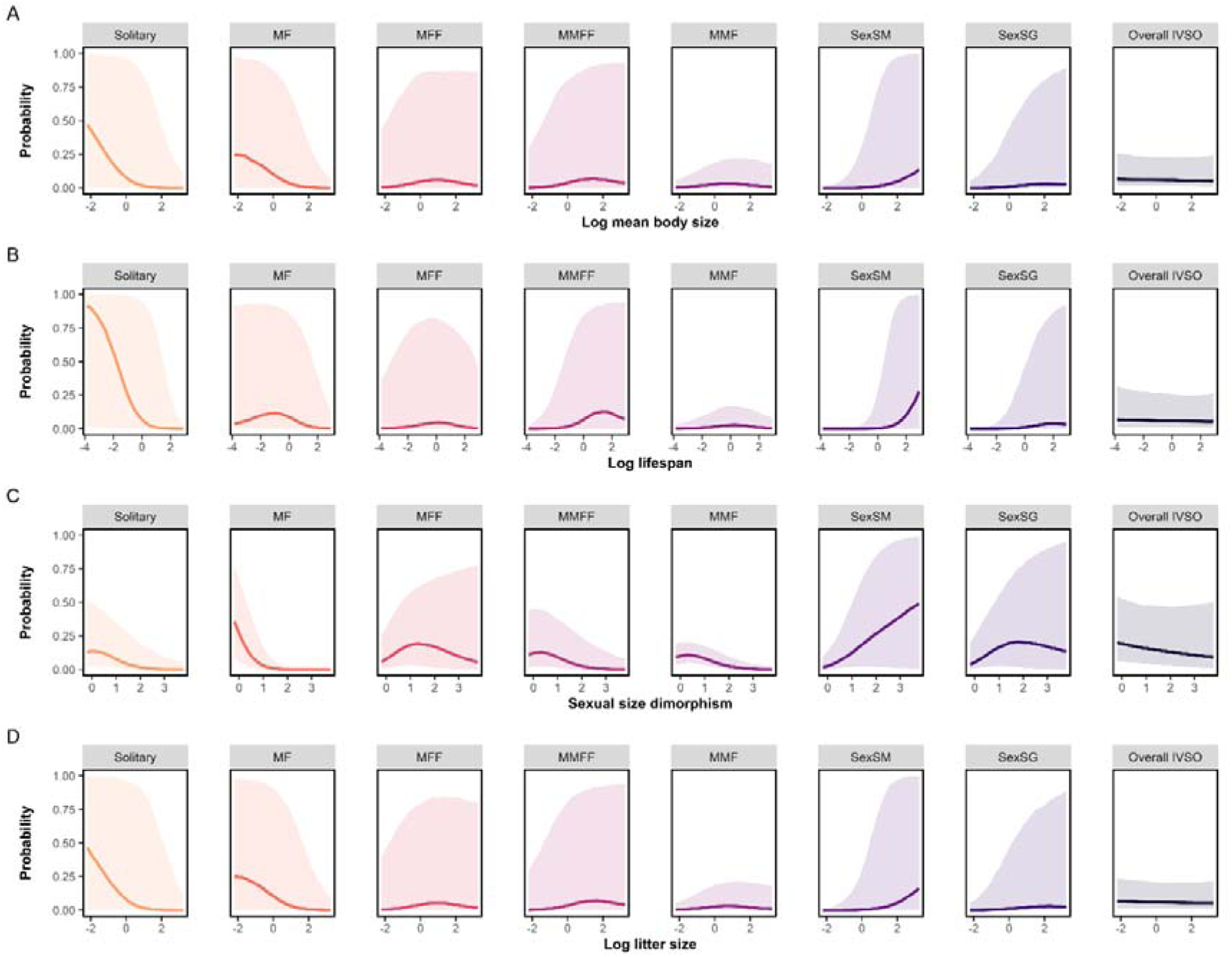
Total effects of continuous predictors on the probability of social organisation across placental mammals, accounting for research effort as well as phylogenetic, species, population-specific effects. Ribbons show 90% CIs. Sexual size dimorphism (SSD) was calculated as the ratio of male to female body size, cantered on zero. For example, an SSD of zero implies no difference between males and females, while a positive SSD implies that males are larger than females.

**Figure 4.**
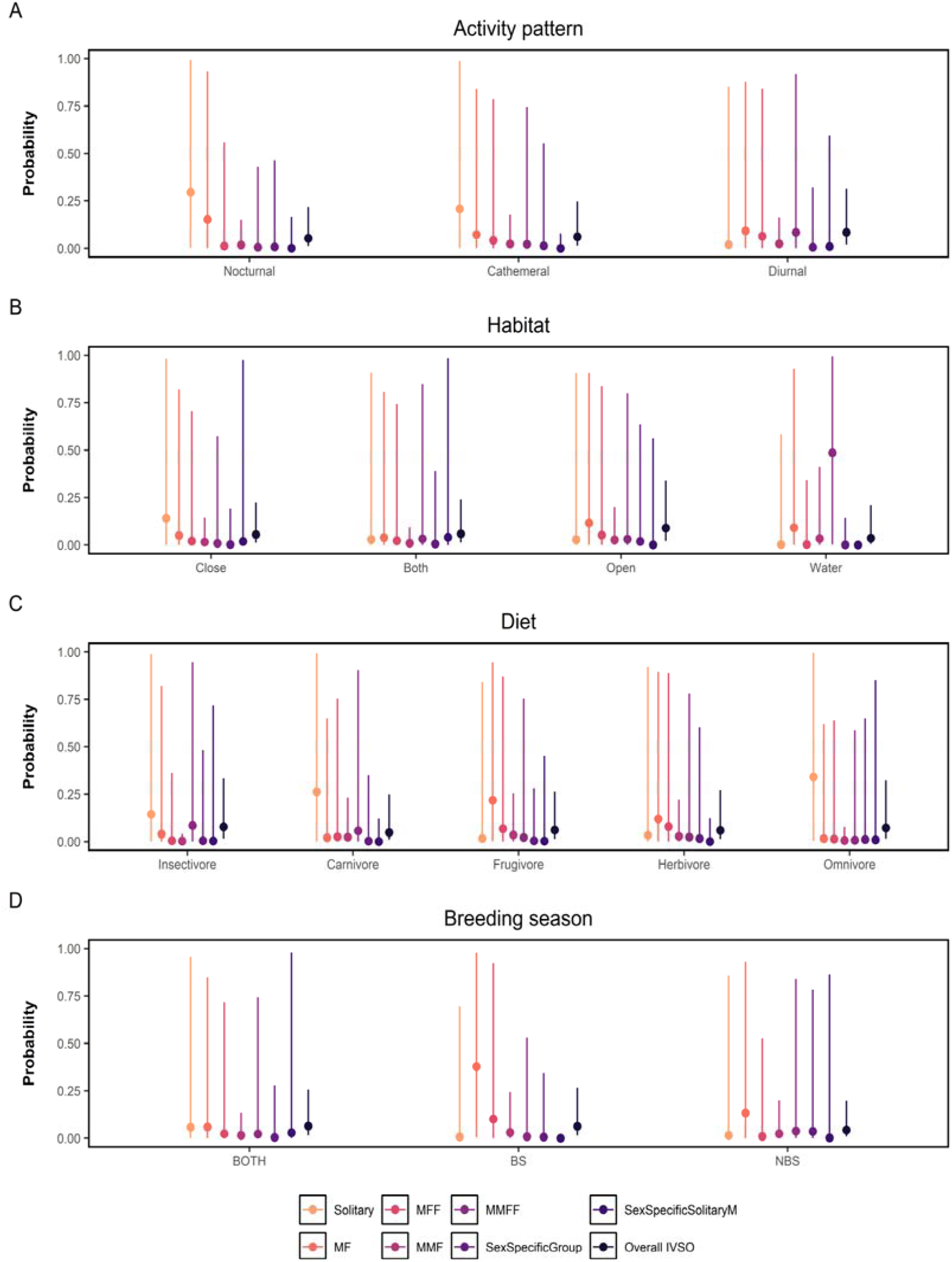
Total effects of categorical predictors on the probability of social organisation across placental mammals, accounting for research effort as well as phylogenetic, species, population-specific effects. Lines show 90% CIs. Breeding season refers to the period during which social behaviour was observed (BS=breeding season, NBS=non-breeding season).

Our database included species with a balance of nocturnal (N=212), diurnal (N=300), and cathemeral activity patterns (N=117). Nocturnal and cathemeral species were most often solitary (nocturnal: 40%, 90% CI 18-99%; cathemeral: 34%, 90% CI 10-99%). Diurnal species were less likely to be solitary than nocturnal species (median Δ in probability = −0.17, 90% CI [−0.74, 8.6×10-4]) and cathemeral species (median Δ in probability = −0.12, 90% CI [−0.64, 3.9×10-4]). Compared to nocturnal species, diurnal species were also more likely to form multi-male multi-female groups (MMFF; median Δ in probability = 0.07, 90% CI [2.4×10-4, 0.62]).

We found that social organization varied with diet. Solitary living was most common for insectivores (33%, CI 0-99%), carnivores (meat and fish-eaters; 39%, CI 0-99%), and omnivores (43%, CI 0-99%; Figure 4C). Insectivores were also somewhat likely to exhibit multi-male multi-female groups (26%, CI 0-95%). In contrast, frugivores and herbivores species were typically either pair-living (frugivore: 34%, CI 0-95%; herbivore: 26%, CI 0-90%) or lived in one-male multi-female groups (frugivore: 21%, CI 0-87%; herbivore: 23%, CI 0-89%). In direct comparisons, frugivores were more likely to be pair-living than both carnivores (median Δ in probability = 0.14, 90% CI [1.9×10-4, 0.68]) and omnivores (median Δ in probability = 0.15, 90% CI [7.2×10-4, 0.70].

We found links between habitat and social organization. Aquatic species were rarely solitary (just 7%, CI 0-58%), and much more likely to form multi-male multi-female groups than species living in open (median Δ in probability compared to open habitats = 0.29, 90% CI [3.5×10-4, 0.85] or closed habitats = 0.37, 90% CI [−1.4×10-3, 0.92]; Figure 4B). Otherwise, species living in closed habitats were often solitary (31%, CI 0-98%), and species living in open habitats were most likely to be pair living (26%, CI 0-91%). There was some, but weaker, evidence that species living in closed habitats were more likely to be solitary than those in open habitats (median Δ in probability = 0.06, 90% CI [−0.02, 0.57]).

Body mass was highly variable across placental mammals in our study, ranging from Whitehead’s woolly bat (*Kerivoula whiteheadi*) at just over 3 grams to the blue whale (*Balaenoptera musculus*) at over 130,000 kg. Solitary and pair-living species were generally smaller, while group-living species were larger (Figure 3A). In a direct comparison, larger species (+ 1SD) were less likely to be solitary than smaller species (−1 SD; median difference (Δ) in probability = −0.13, 90% CI [−0.73, −1.2×10-4]. For example, small species were estimated to have a 46% probability of being solitary (for −2 SD body mass) while the estimate for large species was just 7% (for +2 SD body mass). The uncommon social organisation of group-living females with solitary males was common among larger mammals, such as elephants (median Δ in probability for a –1 SD vs. +1 SD difference in size = 0.01, 90% CI [0, 0.69]). Males were similar in size or larger than females for most mammals in our study, though there were some species where females were larger (e.g., the leopard seal; *Hydrurga leptonyx*). Solitary and pair-living species rarely showed sexual size dimorphism, which was most common in species with one male and multiple females (Figure 3C). More specifically, increased sexual size dimorphism (i.e., males being proportionally larger) was associated with a much lower probability of pair-living (median Δ in probability = −0.66, 90% CI [−0.90, −0.16]), but an increased probability of forming single-male multiple female groups (median Δ in probability = 0.16, 90% CI [0.02, 0.52]), sex-specific social organization (median Δ in probability = 0.15, 90% CI [0.02, 0.53]), or sex-specific female groups with males being solitary (median Δ in probability = 0.12, 90% CI [0.02, 0.48]).

Lifespan was highly variable within our sample ranging from less than one year (e.g., eastern meadow vole*, Microtus pennsylvanicus*) to more than 200 years (bowhead whale; *Balaena mysticetus*). Solitary species typically had short lifespans (Figure 3B), consistent with our finding that longer-living species were less likely to be solitary (median Δ in probability for + 1 SD vs. −1 SD lifespan = −0.18, 90% CI [−0.78, −1.0×10-3]. While some forms of social organisation appeared to be most common at intermediate lifespans (e.g., pair-living, one-male multi-female groups), multi-male multi-female groups, sex-specific groups, and sex-specific solitary male organisation were all more common in long-lived species.

Litter size ranged from small to large for most forms of group-living. Species with larger litters were less likely to be solitary (median Δ in probability for −1 SD vs. +1 SD litter size = −0.12, 90% CI [−0.75, −2.0×10-5]), and less likely to be pair-living (median Δ in probability for −1 SD vs. +1 SD litter size = −0.11, 90% CI [−0.61, 0.02]; Figure 3D). This was unexpected, as generally social species are thought to produce smaller litters of higher-investment offspring ^35^.

Populations observed during the breeding season were most likely to be pair-living (43%, CI 0-98%), and more likely to be pair-living than those observed during both breeding and non-breeding season (median Δ in probability = −0.17, 90% CI [−0.69, −1.1×10-3]). A more even distribution of social organisation was seen in the non-breeding season, or when studies covered both seasons (Figure 4D). We also found that species observed during the breeding season were less likely to exhibit multiple-male one-female groups (MMF) compared to those observed during non-breeding periods (median Δ in probability = −0.08, 90% CI [−0.60, −3.2×10-4]).

### Effects of ecological and life history predictors on IVSO

Intra-population variability in social organization was common (Fig. 2), but did not appear to vary much with ecology or life history (Figure 3, Figure 4).

### Ancestral Social Organisation

Under the assumption that the ancestor of placental mammals was small (approx. 45g), nocturnal, and insectivorous, solitary living was the most likely ancestral social organisation (median probability = 0.45, 90% CI 0.06 – 0.91; Figure 5). There was less, but still substantial, support for the ancestor of placental mammals being pair-living (median probability = 0.26, 90% CI 0.03 – 0.77; Figure 5). This corresponded to a median difference in the probability of the ancestor being pair-living rather than solitary of −0.17, 90% CI [−0.87, 0.68], suggesting that the model could not confidently differentiate whether solitary or pair-living was the main form of social organization estimated for the ancestral population. There was no support for any form of group-living to be the ancestral social organisation, with all median posterior probabilities being below 5% (Figure 5). Inferred ancestral states varied when considering other combinations of diet, activity, and size. However, solitary and pair-living social organization were nearly always found to be the most likely ancestral forms, particularly when assuming nocturnal or cathemeral activity, mean or smaller body size than those in our sample (See Figure S3).

**Figure 5.**
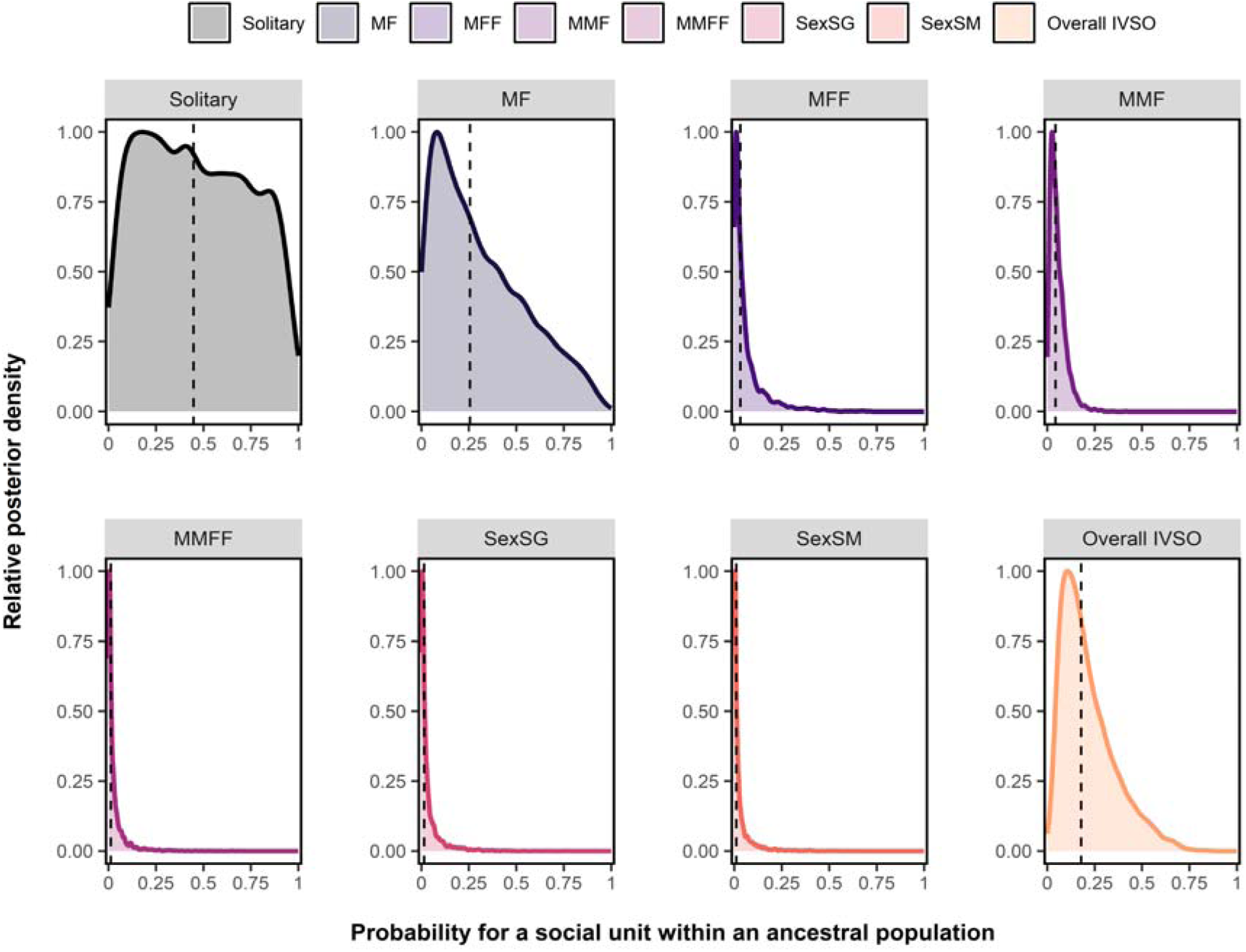
Ecologically informed predictions of social organisation in the ancestor to all placental mammals. Predicted probabilities of a social unit exhibiting each social organisation and some form of IVSO within ancestral populations, assuming expected ecological conditions (nocturnal, insectivore and small), as well as average within-order sampling effort. Scaled posterior densities are shown, with posterior medians indicated by the dotted line.

The probability for overall IVSO (median probability = 0.18, 90% CI 0.05 – 0.5) indicated that approximately 18% of social units in the ancestral population were likely to have deviated from the main form of social organisation. These results were based solely on studies involving more than one social unit (N = 1,001; necessary for IVSO to be assessed), though similar results were found when including all studies (N = 1,276; zero-inflated model: Figure S2, regular model: Figure S3). Therefore, despite the relatively large credible intervals for absolute probabilities, support for moderate IVSO combining solitary and pair-living, rather than group-living, was consistent across analyses and robust to model assumptions.

## Discussion

Consistent with previous studies ^2,4^, we found solitary living to be the most likely ancestral state for placental mammals, though – and in contrast to previous firm assumptions - with a high degree of uncertainty. This uncertainty might be related to the ancestral placental mammal having exhibited intra-specific variation in social organisation (IVSO). Accordingly, and in contrast to previous studies, we also found that a substantial proportion of ancestral placentals were likely to have lived in pairs. This result is consistent with pair-living being inferred as the ancestral social organisation in many placental taxa ^6–8,36^. Our work demonstrates that understanding mammalian social evolution requires consideration of IVSO and datasets based on rigorous empirical data. Unlike earlier work, we did not assume solitary living to be the social organisation for the many non-studied, often small, nocturnal and cryptic species ^10^. Our study highlights the significance of IVSO in mammalian social evolution and shift the focus of research from the origins of pair-living to the origins of group-living.

Our results are not explained by a sampling bias, such as underrepresentation of solitary species or overrepresentation of pair-living species. Our database includes a wide range of solitary species – such as felids, bears, squirrels and several muroid rodents - many of which are well studied due to their conservation relevance or accessibility near research institutions. Extensive field studies have also shown that pair-living is more widespread in small nocturnal mammals than previously believed, particularly in eulipotyphla ^36^ and primates ^8^. Finally, the majority of placental mammals in our database lived in groups containing multiple males and multiple females.

Our finding that the ancestral social organisation of placentals consisted of both solitary and pair-living individuals likely extends to all therian mammals, as a study on marsupials reported a similar pattern ^9^. This suggests that pair-living in mammals originated around 160 million years ago, when marsupials and placentals diverged from their last common ancestor ^37,38^. Our inference based on phylogenetic comparative analysis also agrees with fossil evidence of skeletal aggregations of some cynodont species, indicating that the mammaliform ancestors of mammals might have already been social with parental care 260-230 million years ago ^14,39,40^. Additional support comes from fossil indications that two early mammals were social, the stem-metatherian *Filikomys primaevus* (belonging to the Multituberculata, a side branch of the mammalian tree) ^12^ and the marsupialiformes *Pucadelphys andinus* ^13^. Combining our study and results from palaeontology, there is considerable evidence that ancestral and early mammals were more sociable than generally believed.

We found pair-living to be more common among placental mammals than previously recognised, occurring in 18% of studied populations and across all major orders. Interestingly, the ancestral placental may have shared key ecological and life history traits with passerine birds, which are predominantly pair-living ^41,42^. Fossil evidence suggests the ancestral placentaö was small, insectivorous, and had a central nest (in a burrow) ^11,12,37,43^. A much shorter gestation period than in extant mammals ^11,44^ would have enabled males to participate early in parental care. In passerines, female-female aggression is a main mechanism leading to pair living ^42,45^. As in birds^42,45^, intrasexual aggression, - common in solitary species ^46^ - could have selected for pair-living instead of group-living, especially if females were spatially dispersed. Feeding on widely dispersed insects would have necessitated large home ranges ^47^, making it difficult for males to monopolise multiple females—especially if females were aggressive towards one another. Females’ use of a central nest would have enabled males to monopolize a single partner and increase fitness via mate guarding and possible paternal investment. Additional benefits could have included shared costs of burrow construction and reduced costs of thermoregulation via huddling, a key factor promoting sociality in small mammals ^48^. Thus, pair-living in mammals may have evolved under life history and ecological conditions similar to those in birds, with the added advantage of social thermoregulation.

Our study highlights the emergence of group-living marks a key innovation in mammalian social evolution. As placentals became larger and colonized open habitats, they evolved life-history traits characteristic of modern species ^49^. This fundamentally altered predator–prey dynamics. For small mammals, increased vigilance (e.g. alarm calling ^50^) can be beneficial, but group defence is ineffective against much larger predators. Even a large group of 100 mice would be defenceless against a more than 100 times larger raptor or jackal. Following the Cretaceous–Paleogene mass extinction 65 million years ago, the temporary absence of large predators ^11^ allowed placentals to grow larger and to benefit from group defence. Predator defence is a key benefit of group-living in large mammals and may have driven the evolution of same-sex tolerance, a prerequisite for larger, stable social groups.

Among vertebrates, social groups composed of multiple same-sex adults are rare, seen primarily in a few cooperatively breeding birds ^51^ and some fishes, notably cichlids ^52^. These groups typically represent extended family groups ^53^ that often function as a defence against predators^54^. Large groups are also known from dinosaurs, whose gigantism may have evolved under predation pressure ^54–56^. For mammals, the classic argument for the evolution of groups with multiple males and multiple females is that females grouped together because of predation pressure and resource defense, and that males had to tolerate other males whenever female groups became too large to be monopolized ^27^. On the proximate level, this must have led to the evolution of intra-sexual tolerance in both sexes. Our findings shift the focus of placental social evolution from ancestral pair-living to the phylogenetic novelty of group-living.

While solitary living is ancestral in placentals, it still requires an evolutionary explanation why this form of social organisation persisted for over 100 million years and across diverse environments ^57^. This indicates that solitary living remains the optimal strategy under many conditions. Benefits include reduced risk of disease and parasite transmission, monopolised access to food in territories which can be smaller and are thus more efficiently defended than when shared with conspecifics, and reduced reproductive competition when compared to group-living species with reproductive skew ^58^. For small mammals (and thus for most mammals), reduced visibility to predators is another main benefit.

Life history and ecology were linked to social organisation. Supporting a shared ancestry, solitary and pair-living species shared many traits: small litters, small body size, and absence of male-biased size dimorphism. Group living species were larger and more likely to be diurnal, with a male biased sexual size dimorphism. Only solitary species had a dietary preference, not being specialised on plants as food (frugivorous or herbivorous), while other forms of social organisation spanned the whole range of diets, reflecting the ecological breadth of mammals. In summary, life history traits and ecology were similar between solitary and pair-living species, and among forms of group-living, but different between these two broader categories (Fig. 6).

**Figure 6.**
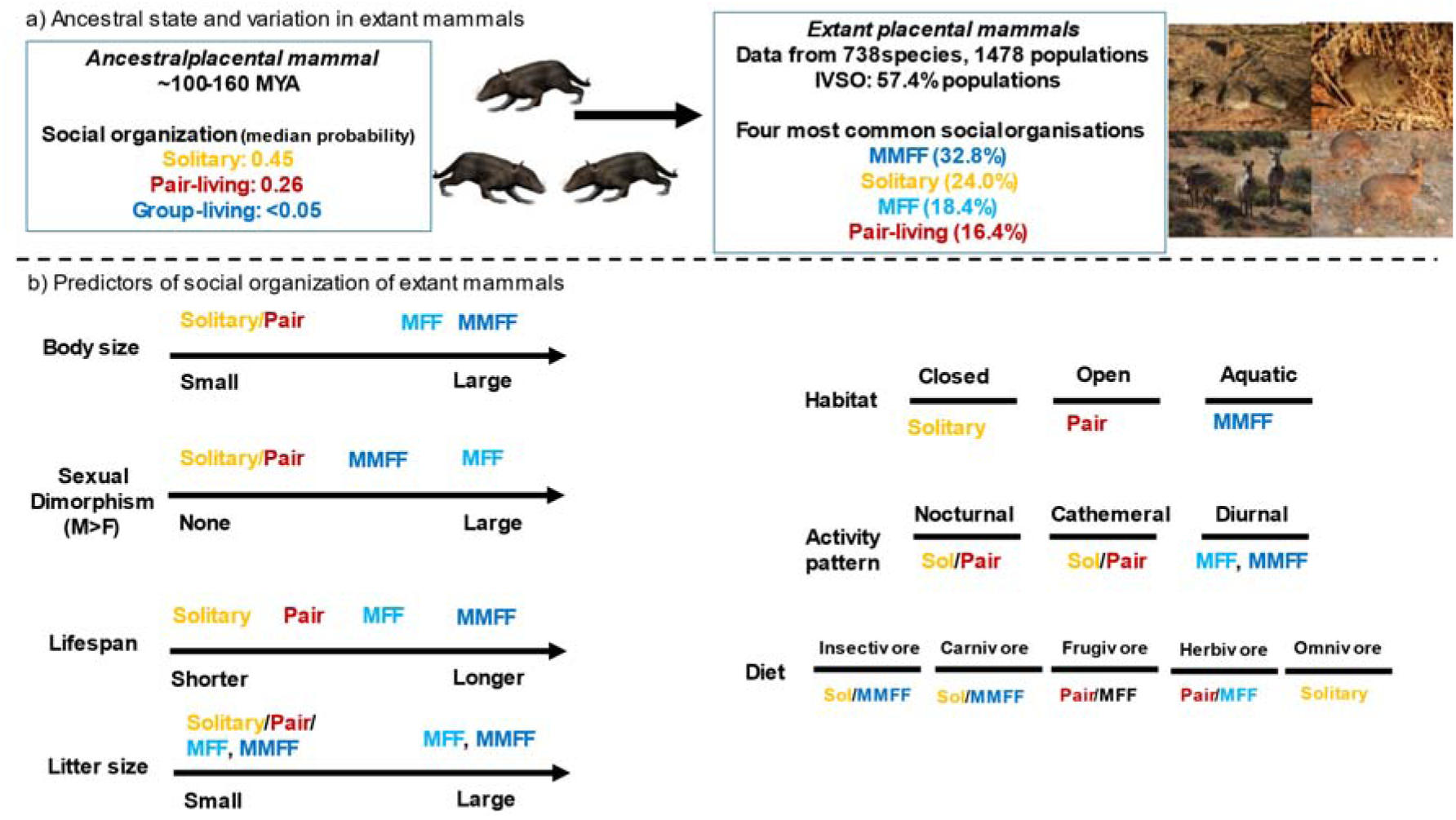
Summary of the main results. a) While the ancestral placental mammal was socially variable, with most being solitary but some living in pairs, in extant placental mammals, multi-male multi-female groups (MMFF) are most common (MFF: one male multi-female groups). b) Ecological and life history factors are related to social organisation in extant mammals, and solitary- and pair living mammals are similar in these factors and different from group-living mammals. Photo credit: Top left, *Juramaia*, an early Eutherian from 160mio years ago. Photo licensed under Creative Commons Attribution-ShareAlike license (CC BY-SA) (© N. Tamura). Top right: Striped mice (*Rhabdomys pumilio*) living in multi-male, multi female groups, solitary bush Karoo rat (*Otomys unisucatus*), Hartmann’s mountain zebra (*Equus zebra hartmannae*) living in single male multi-female groups, and pair living Kirk’s dik-dik (*Madoqua kirkii*). All photos © C. Schradin.

Our study focussed only on social organisation, not on social structure, i.e. social interactions within pairs and groups. For example, whether pair-bonding – present in many but not all pair-living mammals ^18^-is ancestral or evolved multiple times via distinct physiological mechanism remains an open question. We therefore advocate for future research on social tolerance, bonding mechanisms, and the costs and benefits of solitary living.

In summary, our results suggest that the ancestor of all placentals was primarily solitary living, but that pair-living as an alternative form of social organisation occurred in a significant part of the population. The substantial presence of pair-living in early placentals may explain its ancestral status in multiple orders across three of the four placental superorders - Afrortheria (elephant-shrews^6^), the Laurasiatheria (artiodactyla ^7^) and the Euarchontoglires (primates ^8^). The novelty in placental social evolution was the occurrence of group-living, especially multi-male multi-female groups, which could only arise with the evolution of intra-sexual tolerance.

## Methods

### Collecting measures of social organization across mammals

The 5390 species of placental mammals listed in the IUCN (International Union for Conservation of Nature) database (2019) were considered. We used a recent and comprehensive phylogeny of mammals to measure the historical relationships between these species (Vertlife Project ^59^). We then conducted literature searches on field studies reporting the social organisation for each of these mammalian species. The social organisation of a species is described by the size, the sexual composition and the spatio-temporal cohesion of the group^15^. Our analysis focuses on group composition as a measure of social organisation that can be robustly identified in field studies for many species. Similar to Olivier et al. ^8^, we considered several categories of social organisation (for definitions see Table S1): solitary, pairs (MF), one male with multiple females (MFF), multiple males with one female (MMF), groups of multiple males and multiple females (MMFF), sex-specific groups, sex specific solitary male (females formed groups while males were solitary), sex specific solitary female (males formed groups while females were solitary, but this was never found).

For each species, we searched the Latin name of the species and the term “social” (*e.g.* “*Rhabdomys pumilio* AND social”) using Web of Science (Thomson Reuters) and Google Scholar. Searches occurred between December 2019 and September 2022. For Web of Science searches, we only considered results within the subsections of “behavioral science”, “environmental science/ecology”, and “zoology”. If no study was found, we conducted additional searches based on Latin name only. For species where the Latin name had changed over time, we conducted separate searches for each version of the name.

The target of our search was primary peer-reviewed literature documenting field studies of social organisation; thus, we excluded review articles. We also excluded any studies based on animals in captivity or enclosures smaller than 1000 ha, as their group composition and social behaviour can be influenced by their captive environment. Studies that included manipulation of individuals (*e.g.* adding or removing individuals), groups or resources were discarded. We also excluded studies that did not report the sex or that could not determine if individuals were adults or subadults, following ^60^. In the remaining articles, data for group composition were extracted from the methods, results, figures and tables section, avoiding any interpretations from the authors in the discussion.

It is typical for studies of social organisation to report group composition for a number of social units (e.g., observations of groups of animals). As categories of social organisation (MMF, MMFF, etc.) may vary across groups within a population, we recorded the frequency of social units of each category for each study. As we were interested in the sex composition of adults, we did not consider juveniles, subadults and pups. We used data on home range sizes and spatial overlap, for example from radio-tracking studies, to determine social organisation. However, overlapping home ranges alone does not indicate social association, or shared membership in a group. Only when individuals shared the sleeping sites or when the home ranges of several groups were clearly distinct from each other, were they considered to be pairs or groups, and included in our analysis.

We recorded social organisation of observed social units using specific criteria ^60^, as used in a previous study ^8^ and explained in detail in the supplementary materials and methods S2. Solitary was only recorded when both males and females were observed to be solitary, and we excluded any dispersers, because they occur in nearly all species. We also did not consider data from aggregations for mating or at temporary food sources.

Data were collected at the population level by recording the total number of papers reporting a given social organisation in a population, i.e. a particular field site. When the same observed individuals and their social units were included in more than one published paper, we considered only the most precise paper, *e.g.* papers where the precise number of social organisations and/or the sex of individuals was described, to prevent the same social unit from being counted several times. The total number of studies reporting social organisation per population was then recorded in the database to control for any effects of research effort, as this influences the probability to find evidence for IVSO. In addition, we recorded whether the study took place during the breeding season, during the non-breeding season, or throughout the year.

Some populations in our database exhibited “multilevel” social organisation ^61^, where animals associate within a hierarchy of units: e.g., several MFF groups constitute a clan, several clans form a band, etc. In these cases, we only recorded the composition of the core (most basic) level of groups defined in the study. Many species exhibited fission-fusion social behaviour, temporarily forming groups varying in size, composition, and duration of association. In these instances, we recorded all forms of group composition observed, as this indicated that individuals of this population could exhibit multiple forms of social organization within their society over time. When sex was known, studies reporting groups of adult siblings or groups of a pair of breeding adults and their adult offspring were reported as multi-male multi-female groups rather than assuming kinships relations, as most studies did not represent genetic data on relatedness. For example, callitrichid primate MMFF groups were previously often believed to represent one breeding pair and adult offspring, but later genetic studies indicated this often to be incorrect, with more than one male and/or female breeding per group ^62^.

The database and R scripts will be published ^63^ and are available for review here: https://data.indores.fr:443/privateurl.xhtml?token=917175ed-1794-4ad8-8eda-78e8248f3f4d.

### Calculating intraspecific variation in social organization (IVSO)

IVSO was defined as the detection of multiple forms of social organization in a given population. However, the following cases were not regarded as IVSO: dispersing individuals (solitary individuals of one sex only) or alternative reproductive tactics. Environmental disruption such as the death of a dominant breeder or predation of group-members can also change the social organization of a unit ^64^, but these changes do not reflect the evolved behavioral plasticity we want to explain. Thus, such environmental disruption events were not considered in our database.

In contrast to previous studies that considered IVSO as a categorical variable ^5–7,9,36,65^ we treated it as a continuous variable, following prior work on primate social orgainzation^8^. We calculated IVSO at the population level as the proportion of social units deviating from the most frequently observed social organization (‘main social organization’) within a population. Thus, we conceptualised and measured IVSO as a distinct trait capturing the degree to which a population’s social organization was dominated by a single category ^8^. This variability, or lack thereof, may itself co-evolve with the composition and frequency of specific social organizations within a population. Specifically, we conceived of IVSO as the proportion of observed social units deviating from the most common form of social organization within a population. This can be expressed as:

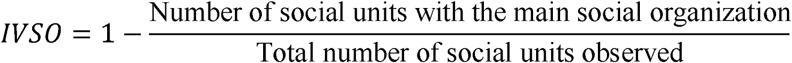

This quantitative approach avoided the use of arbitrary thresholds for categorizing the presence or absence of IVSO, retaining the continuous information provided by previous literature, and allowed us to consider how the evolution of specific forms of social organization and the overall degree of IVSO are related across species and populations.

### Collecting measures of key predictors of social organization

We included the following predictors in our Bayesian model to account for potential social, ecological, life-history and methodological causes of variation in social organization: habitat type, body mass, diet, activity pattern, longevity, litter size, sexual dimorphism, seasonality and number of studies per population (Table S2).

Habitat type was recorded from the primary literature and categorized using IUCN classification. Given the large number of habitat categories that exist (15 habitats), we then classified them into open, closed, open and closed (both) or water. For the other predictors (body mass, diet, activity pattern, litter size, sexual dimorphism) we used three different databases to obtain the most accurate information for each species, the Handbooks of the Mammals of the World ^66^, the online database Pantheria ^67^ and the online database AnAge ^68^. First, we compared the three databases to see if their information was similar (the correlations between the three databases were between 0.92 and 0.99). The most precise information (*e.g.* data for males, females, ranges and mean) was available in Handbook Mammals of the World, which we therefore privileged. If no information was found in the Handbook Mammals of the World, we then consulted the online database Pantheria and again if no information was found, we used AnAge.

### Modelling the predictors of social organization across species

To determine the ancestral social organization of all mammals, IVSO across mammals and the different ecological and life history factors associated, we developed multilevel phylogenetic models following the analytic framework developed in prior work on IVSO among primates^8^. All analyses were conducted within a flexible Bayesian framework to account for non-Gaussian outcome measures, repeated measures of the same species, as well as uncertainty in phylogenetic relationships across taxa ^69,70^. Main social organization was modelled as a multinomial response variable, while IVSO was treated as a binomial response. Multi-response models were estimated to assess phylogenetic and ecological effects and to conduct robust ancestral state reconstruction of the main social organization and the degree of IVSO expected in an ancestral mammalian population. We used weakly regularizing priors to have more realistic assumptions about the estimators and to reduce the risks of inferential bias caused by multiple testing and measurement and sampling error ^70,71^. Effects are reported either as full posteriors or as medians of the posterior with 90% credible intervals (CIs).

For each predictor, we fit two generalized multilevel phylogenetic models (see ^8^ for further details) The first explained the probability of a population exhibiting each type of social organization with a multinomial distribution. For population *p* for species *s*, the multinomial model predicted the number of units observed in category *i* as a function of the total number of units observed *n_PS_* and a vector of effects, *θ_PS_*, expressing the relative probability of a population belonging to each category, compared to the reference category, which in this case was multi-male groups with single female (MMF) social organization. This was chosen as the reference category as it was expected to be relatively rare, and unlikely to be the ancestral state of mammalian social organization. However, we focus on absolute probabilities rather than these specific comparisons estimated by the model when reporting the results.

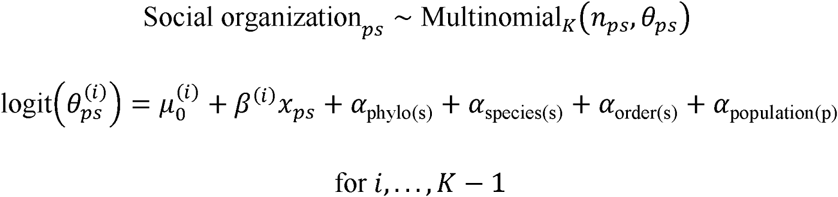

Here, *i* indexes the categories of social organization (excepting the reference category). Intercepts are represented by *µ*_0_, while *β* captures fixed effects of key predictors included in the model (e.g., diet, research effort, lifespan, etc.). Also included are varying intercept effects *α* capturing variation resulting from phylogenetic distance, as well as specific to populations, species, and orders. Phylogenetic effects were fit using a Brownian motion model which assumes that the expected covariance between species should decline linearly with phylogenetic distance.

We used a similar approach to explain variability in social organization (IVSO). We used a binomial model whereby the number of social units differing from the most common category of social organization were expressed as a function of total social units observed for a given population (*n_PS_*) and the expected rate of deviation, *τ_PS_*. Otherwise, the same varying effects structure was used for these models as well. While the binomial approach accounts for the total number of social units, deviation from the most common form is not possible when just a single unit was observed. For this reason, we re-ran IVSO models while excluding these populations to assess if results were similar. Sample sizes for these models varied depending on the availability of measures of predictors across species (Summarized in Table S5).

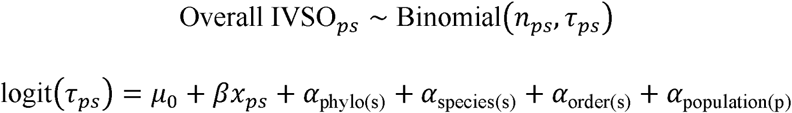

### Inferring ancestral social organization

Comparative phylogenetic models can be used to infer the ancestral states of biological traits, including behaviour. While ancestral states can be inferred from the intercept of comparative phylogenetic models (i.e., prior to the influence of species-specific or phylogenetic effects) ^69,72^, blending in data on fossil species allowed us to minimize potential biases resulting from unbalanced samples across the 100 - 160 million year of placental evolution ^37,73–76^. Recent studies have revealed that the ancestor of all placentals was small with a body mass of below 50g, nocturnal and insectivore ^37,43^. Body size (=−1.9SD), diet (=insectivore) and activity pattern (=nocturnal) were therefore used to make more informed inferences about ancestral states, following^8^, whereas habitat, litter size, sexual dimorphism and longevity were excluded from this model as there was no information available in the literature about a priori expectations on those variables for the ancestor. As robustness checks, we also report the estimated ancestral social organization for all possible combinations of a range of body sizes (−2 SD, −1 SD, mean, + 1SD, +2 SD), all diet categories, and all activity patterns in our analysis. We also adjusted for differences in research effort across orders.

All analyses were conducted in R^77^, using the *brms* package to fit models ^78^. We also used the *targets* package to organize analyses ^79^. The R scripts will be published ^63^ and are available for review here: https://data.indores.fr:443/privateurl.xhtml?token=917175ed-1794-4ad8-8eda-78e8248f3f4d.

### Text editing

All text was first written and edited by us. Especially text from the introduction and discussion was then also edited via ChatGPT using the prompt “Act as a scientific editor. Change the text below to read better. Also correct any grammatical errors. Keep it short and precise. Suggest all changes and mark them for me to review.” Suggested changes were then reviewed and incorporated when found to improve the manuscript. The new version was then again read and edited by all co-authors.

## Acknowledgments

Computational resources were provided in part by ACENET (ace-net.ca) and the Digital Research Alliance of Canada (alliancecan.ca).

## Author contributions

Conceptualization: CS, CAO, LH, AVJ

Methodology: SW, JSM, AVJ, CAO, CS, LM

Investigation: SW, CAO, LM, JQ, AVJ, CS

Visualization: SW, LH, CS, JSM, AVJ

Funding acquisition: CS

Project administration: CS Supervision: CS, AVJ

Writing – original draft: CS, SW, CAO

Writing – review & editing: CS, SFW, LH, LM, CAO, JSM, AVJ.

## Data and materials availability

^63^https://data.indores.fr:443/privateurl.xhtml?token=917175ed-1794-4ad8-8eda-78e8248f3f4d

## Supplementary Materials

### Supplementary Figures

**Figure S1.**
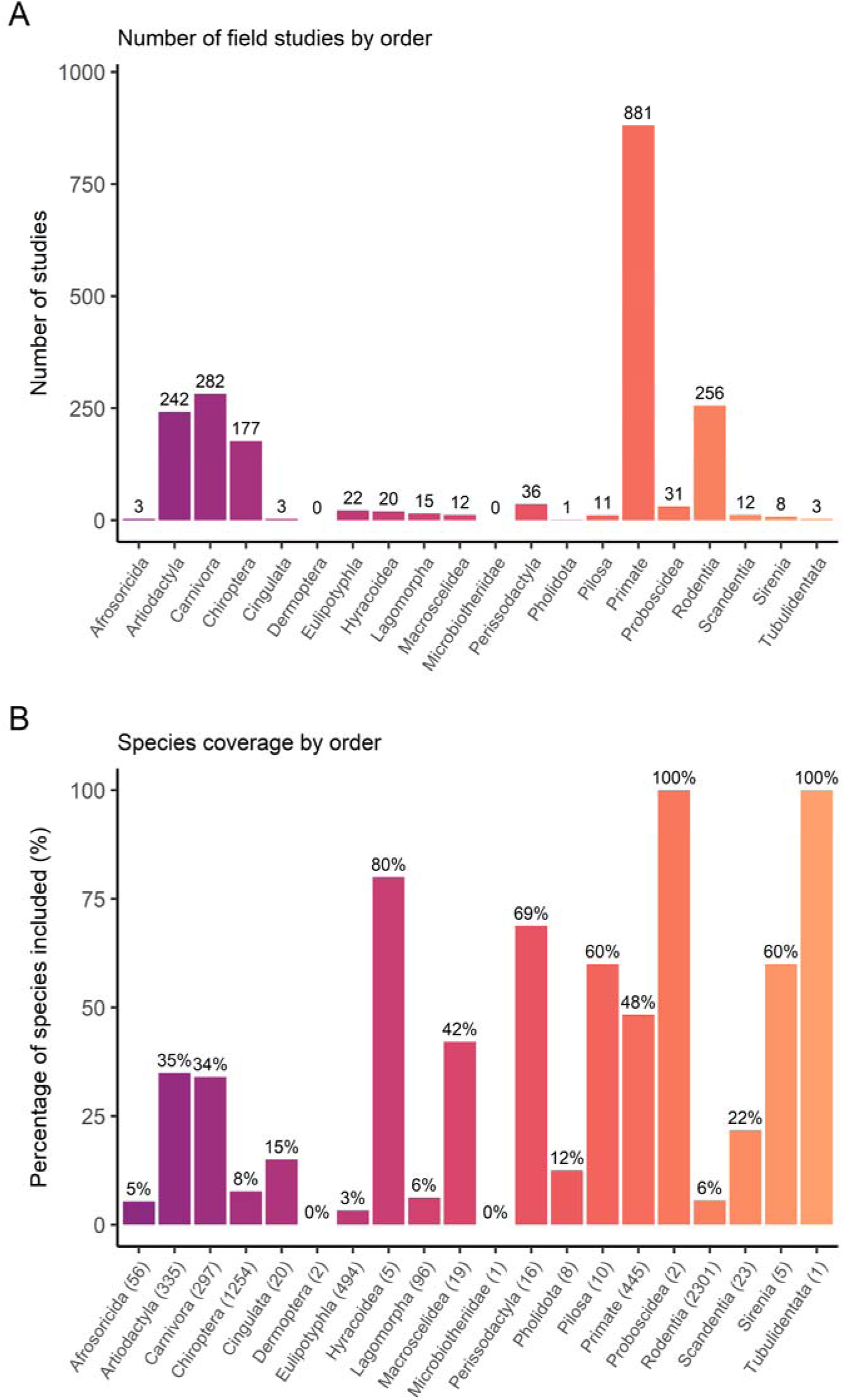
In proportion to the number of extant species, we found that primates were the mammalian order with the most field studies in the primary literature (881 studies) followed by carnivores (282 studies), artiodactyls (242 studies), rodents (256 studies) and bats (177 studies; Figure 1A). A) Number of field studies on social organization for each order of placental mammals (number of species are indicated on each bar). B) Percentage of species with information on social organization for each order. The percentages are indicated on each bar, while number of total species per order are indicated on the x axis. The total number of species per order were based on the most recent version of a database provided by the American Society of Mammalogists Database ^1^.

**Figure S2.**
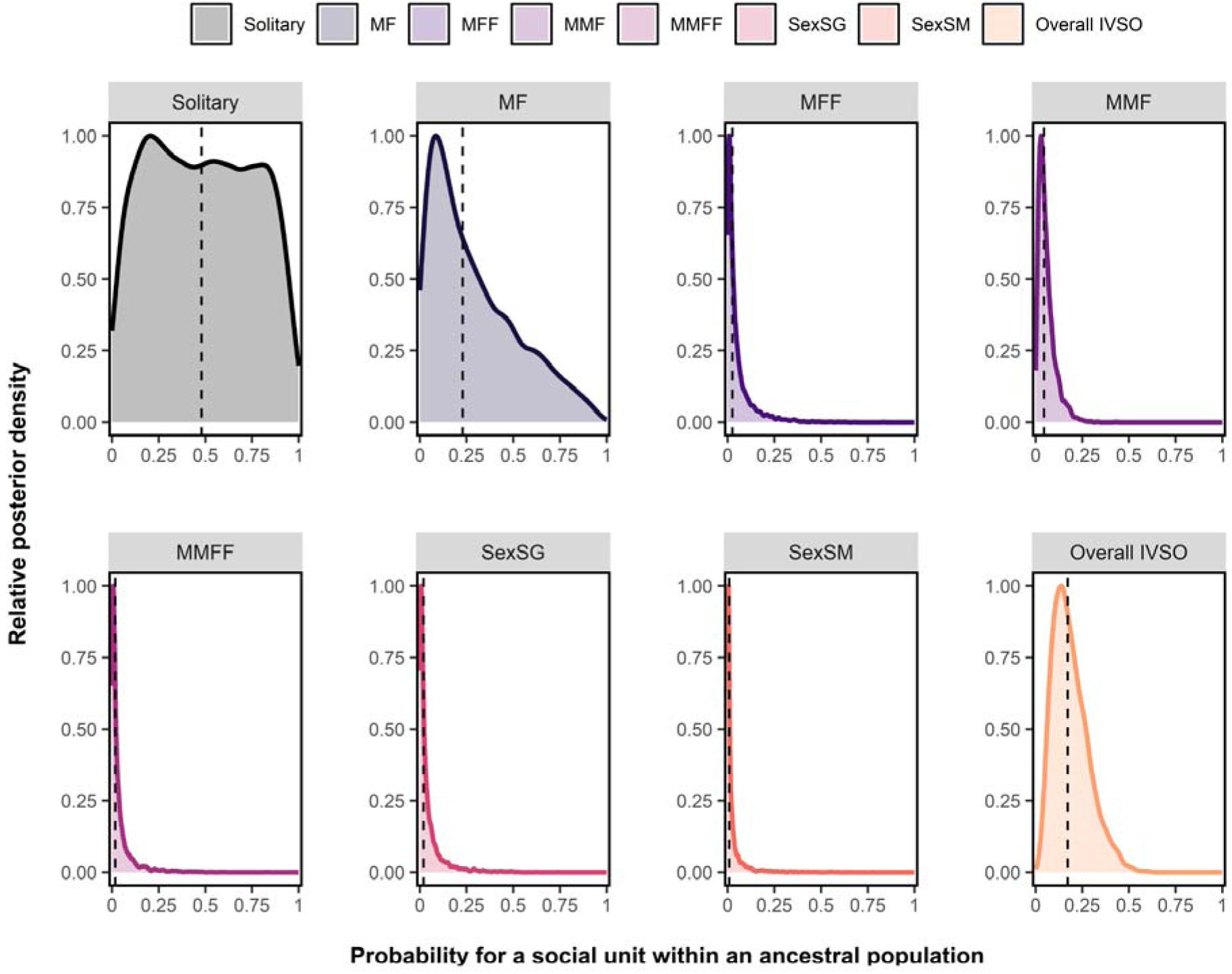
Ecologically informed predictions of social organization in the ancestor to all placental mammals. Here we included also studies that reported data from only one social unit. We used a zero-inflated model structure to account for excess zeros. Predicted probabilities of a social unit exhibiting each social organization and some form of IVSO within ancestral populations, assuming the ecological conditions (nocturnal, insectivore and small), as well as average within-order sampling effort. Scaled posterior densities are shown, with posterior medians indicated by the dotted line. For abbreviations see legend Figure 1 main text.

**Figure S3.**
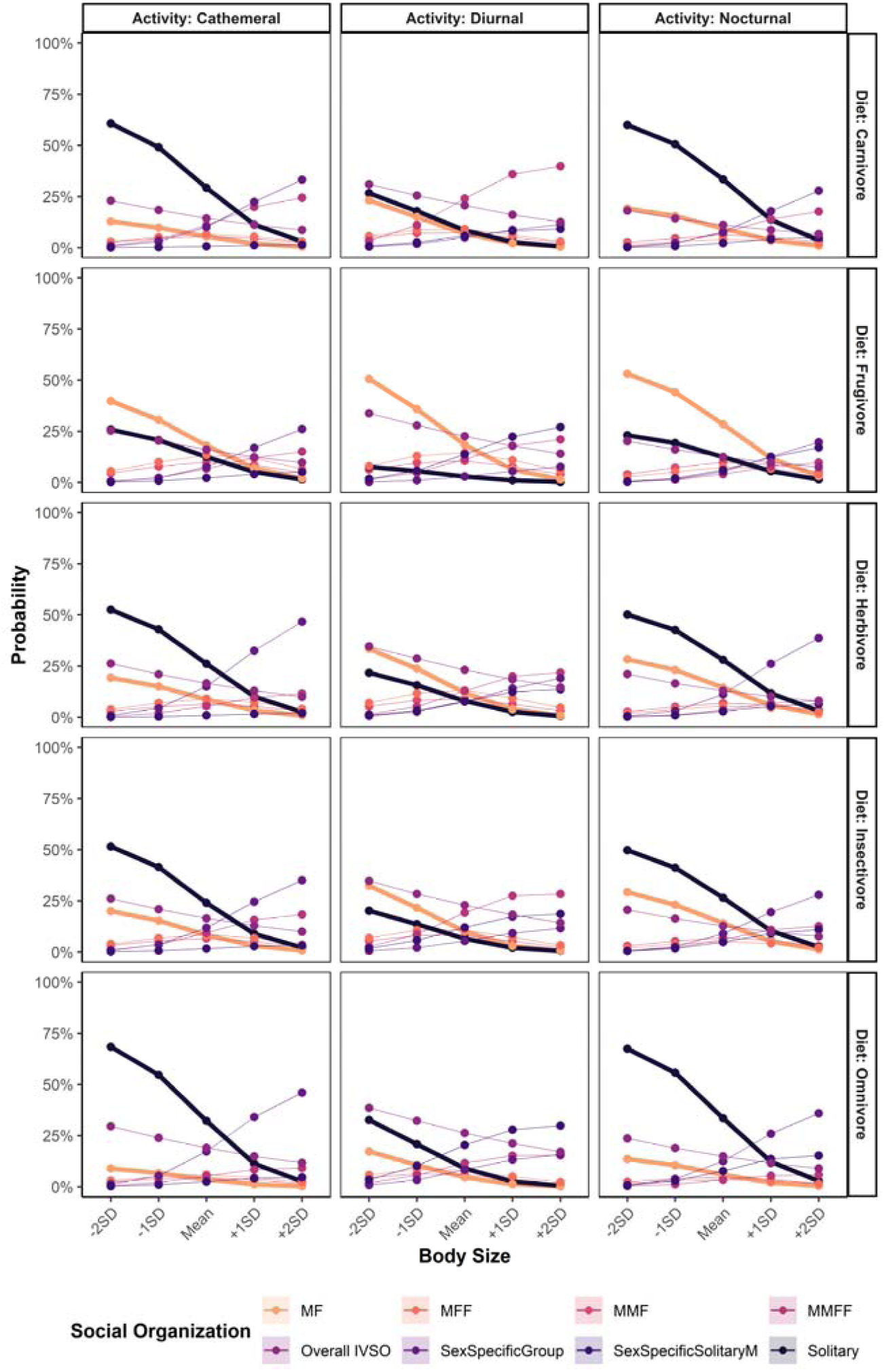
Probabilities of ancestral social organization based on various combinations of diet, activity pattern, and body size. Points show the medians of posteriors of the absolute probability of the ancestor of placental mammals forming each type of social organization. Solitary and pair-living lines have been bolded for easier comparison. While solitary and pair-living social organization were often most likely to be inferred for each hypothetical ancestor, inferences varied, particularly if the ancestor was larger than extant mammals, was diurnal, and/or had a frugivorous diet. For abbreviations see legend Figure 1 main text.

### Supplementary Materials and Methods S1

We tested whether ecological and life history factors were associated with specific forms of social organization. Specifically, we predicted that diet influences sociality: species feeding on non-sharable food (insects and other animals) are more likely to be solitary, while species feeding on sharable food (e.g., plants, fruits) are more likely to be group-living. Species in open habitats were predicted to form groups to reduce predation risk. We further predicted sexual dimorphism to be lowest in pair-living species and largest in group-living species. We also tested effects of body mass, longevity and activity pattern on sociality predicting that small, nocturnal species are more likely to be solitarily, while long-lived diurnal species are more likely group-living. Finally, we tested for factors associated with IVSO. Successive generations of short-lived species and species with only few breeding attempts per generation were predicted to be more variable in their social organisation to reproduce under several environments which differ considerably but predictably from the environment of other generations. Alternatively, long-lived species with multiple breeding attempts may encounter different environments during their lifetime, making IVSO adaptive.

### Supplementary Materials and Methods S2

Only instances in which both sexes were observed to be solitary were counted as “solitary” social organization, so as not to confuse temporarily dispersing animals which may ultimately join others. Whenever individuals were explicitly reported to be dispersing, they were not considered in the recording of social organisation. Therefore, to record one social unit of solitary living, at least one solitary male and one solitary female were needed. For example, when 4 solitary males and 2 solitary females were reported, we recorded 2 solitary social units (the minimum between sexes). The same procedure was applied when classifying sex-specific groups. More specifically, we only recorded sex-specific groups as a form of social organisation when at least one group of males and one group of females had been observed. For example, when 10 groups of males and 4 groups of females were reported, we recorded 4 sex-specific groups social units. Similarly, observations of at least one solitary male and a group of females were required to have a sex-specific solitary male social unit. Though we would apply the same logic to sex-specific solitary female social organisation, we did not record any instances of this category in our database.

We did not consider animals to form pairs or groups if they were reported as only being together due to mating or foraging aggregations (e.g., solitary individuals attracted to the same clumped food source). Considering the huge variation in life history between mammalian species as well as duration of field studies (from weeks to decades), it was not possible to define a time period animals had to spend together to be recorded as pairs/groups that was suitable for all species and studies; instead, we relied on the assessment of the field researchers reporting animals as living in pairs/groups or not in the primary literature. In cases where it was not clear from the presented data whether individuals observed together formed pairs/groups outside of the context of mating, we excluded the study in question from the database (see also ^2^).

We considered all social units reported in papers to calculate intra-specific variation as the deviation from the main form of social organisation. For example, if for 3 species 10 social units were observed and pair-living was the main form of social organisation, within species A all individuals living in pairs, in species B 75% and in species C 60%, then IVSO was calculated as 0.00, 0.25 and 0.40, respectively.

Whenever we were unsure which social organisation was reported in the primary literature and whether any possible deviation from the main form of social organisation was reported or not, we did not include the study in our database. For example, studies which did not report any data on social organisation but only mentioned the assumed main form of social organisation in the introduction and discussion, and not whether there was any deviation from it (IVSO), were excluded. Thus, only data, not general statements entered our database.

### Supplementary Tables

**Table S1.** Definitions and sources of predictors used in analysis.

| Predictors | Definition | Database <sup>3-5</sup> |
| --- | --- | --- |
| Habitat type | The type of habitat depending on whether it is open, closed, both, or water (i.e., exclusively aquatic).<br>Open: grassland, shrubland, rocky areas, savannah, desert.<br>Closed: forest, wetland, cave, woodland.<br>Both: when a population lived in an open habitat (*e.g.* grassland) and in a closed habitat (*e.g.* forest).<br>Water: sea, river, oceans, lake. | Based on our own primary literature search. |
| Body mass | Mean body mass (data of males and females combined). | Handbook Mammals of the World, Pantheria, AnAge. |
| Diet | Insectivore, carnivore, frugivore, herbivore, or omnivore. | Handbook Mammals of the World. |
| Activity pattern | Diurnal, nocturnal, or cathemeral. | Handbook Mammals of the World, Pantheria, AnAge. |
| Longevity | Mean longevity. | Pantheria, AnAge |
| Litter size | Mean litter size. | Handbook Mammals of the World, Pantheria, AnAge. |
| Sexual size dimorphism | Male body mass divided by female body mass. | Handbook Mammals of the World, Pantheria, AnAge. |
| Breeding season | Whether observations of social units occurred during the population's breeding season, non-breeding season, or both. | Based on our own primary literature search. |

**Table S2.** Forms of social organization recorded in our study and their definition. “Stable” refers to pairs or groups of the same individuals being repeatedly observed, also outside the context of mating. Note that we did not find any instances where mammalian social organization was classified as sex-specific solitary female (i.e., groups of males and solitary females).

| Social organization | Definition |
| --- | --- |
| Solitary | Adult males and females forage independently and only meet for mating (do not stay together for periods longer than courtship and mating) and are otherwise alone. All identified dispersers were excluded. Cases where only one sex occurred alone were considered as potential dispersal or alternative reproductive tactics and were not accounted as a solitary social organization. |
| Pair-living (MF) | Repeated observations of one individual adult male and one individual adult female together, with or without dependent offspring. The home ranges of a pair overlap with each other to a great extent (>>80%) but not with others. |
| Groups of one male and multiple females (MFF) | Observations of stable groups with one adult male and multiple adult females with or without dependent offspring. |
| Groups of multiple males and one female (MMF) | Observations of stable groups with multiple adult males and one adult female with or without dependent offspring. |
| Groups of multiple males and multiple females (MMFF) | Observations of stable groups with multiple adult males and multiple adult females with or without dependent offspring. |
| Sex-specific groups | Both multiple adult male groups and multiple adult female groups occur in the population. Cases where only one sex formed-groups as an alternative reproductive tactic or non-reproductive tactic (bachelor groups) were not considered as a sex-specific social organization. |
| Sex-specific solitary: male | Solitary adult males and groups of adult females occur in the population. |
| Sex-specific solitary: female | Solitary adult females and groups of adult males occur in the population. |

**Table S3.**
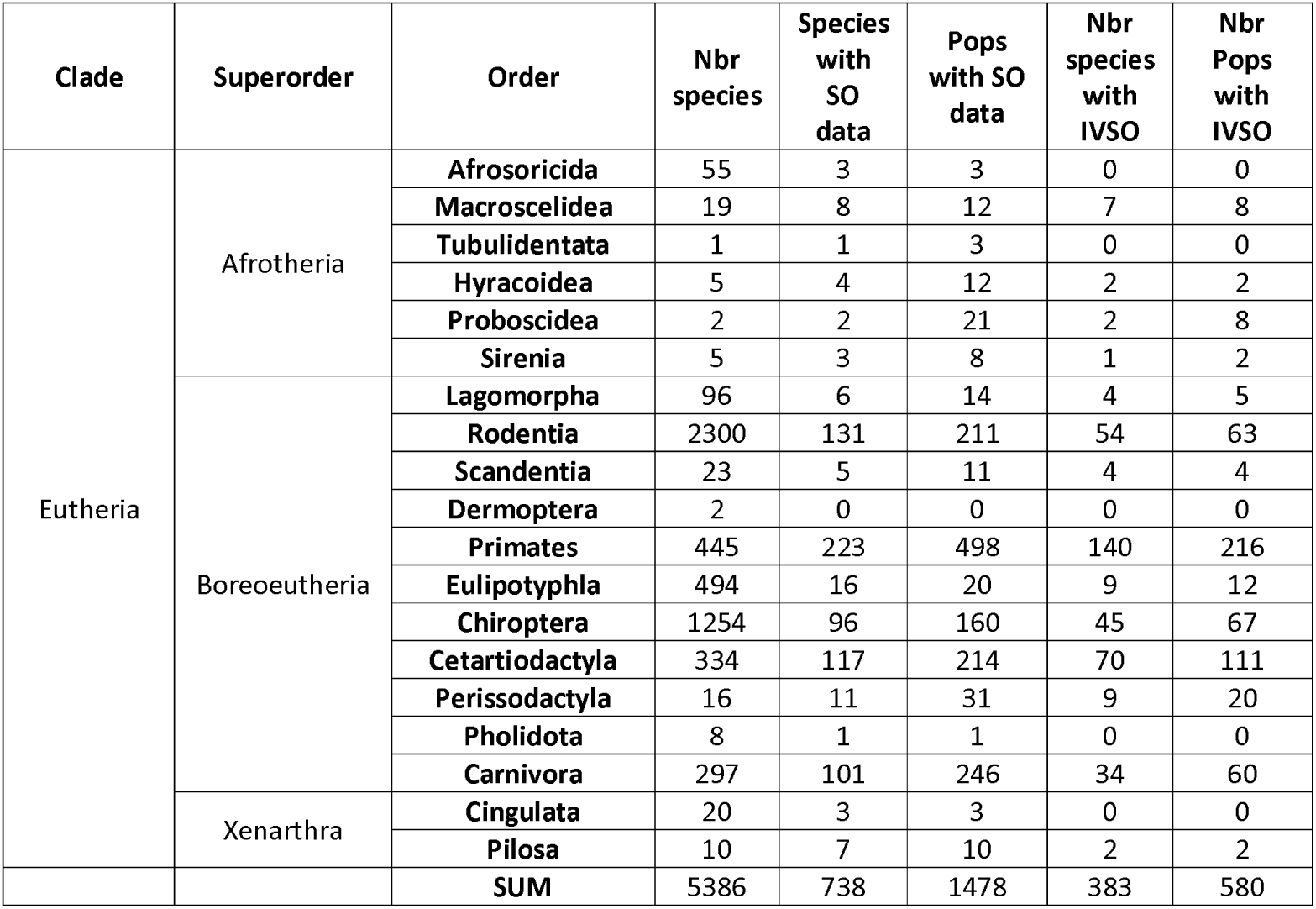
The total number of species for each mammalian order is indicated as well as the number of species and populations with information on their social organization. The number of species and populations with more than one form of social organization is also provided.

**Table S4.**
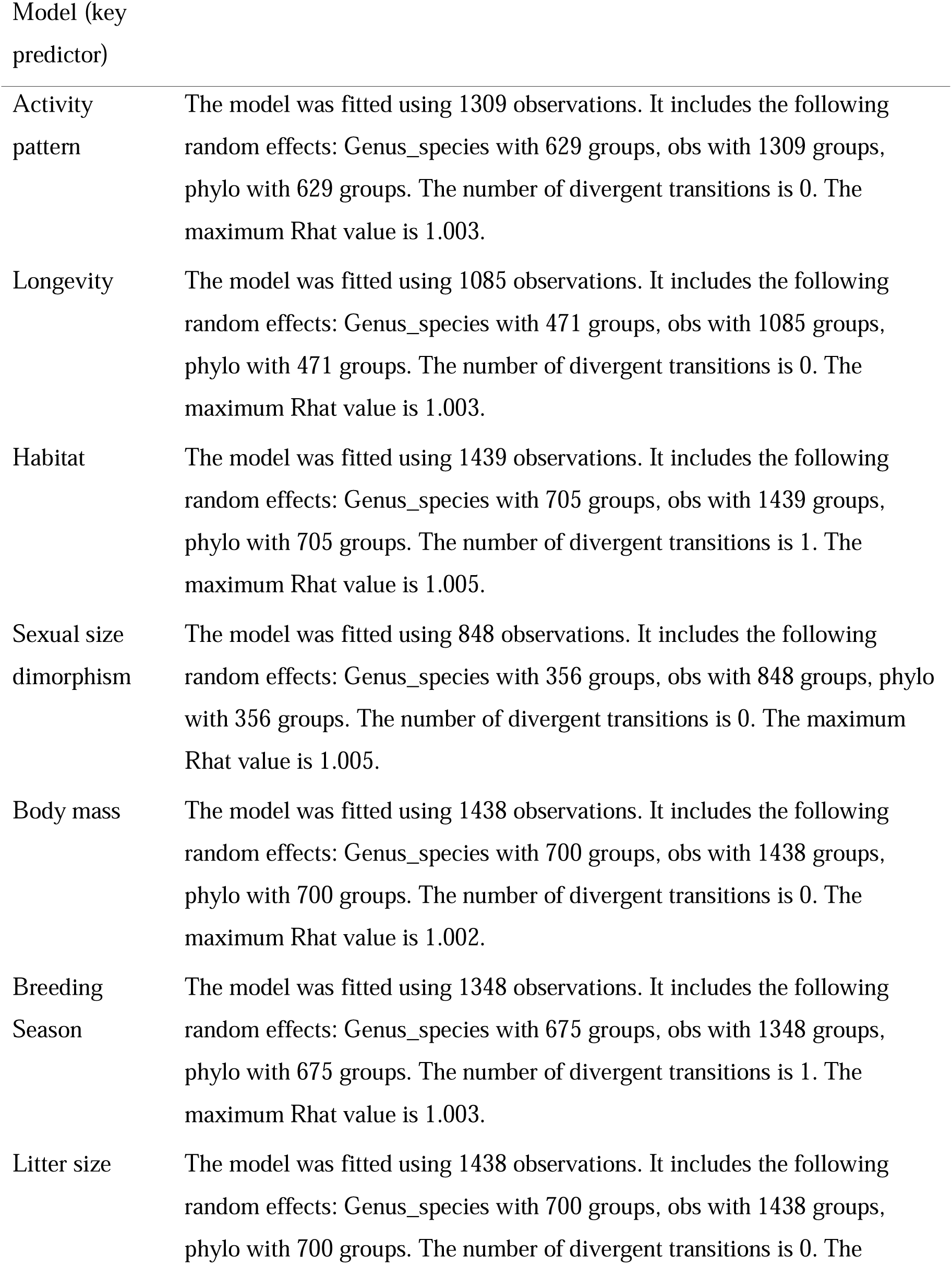
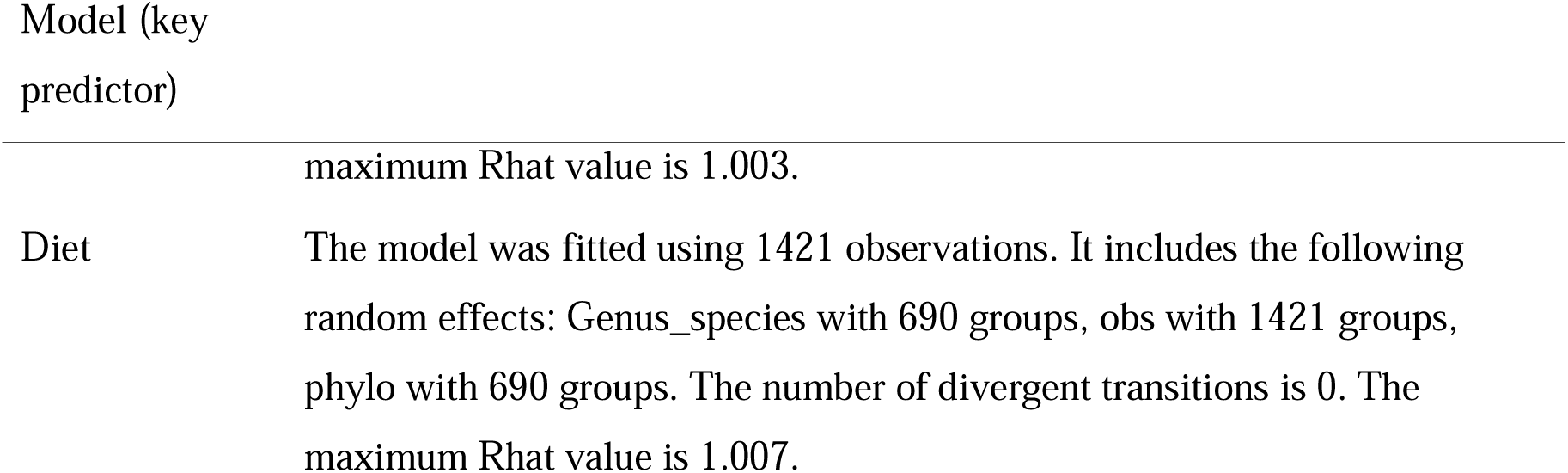
Summary of sample sizes for Bayesian phylogenetic GLMMs. Here “obs” refers to populations while “phylo” and “Genus_species” refer to phylogenetic and additional species-specific varying effects.

| Model (key predictor) |  |
| --- | --- |
| Activity pattern | The model was fitted using 1309 observations. It includes the following random effects: Genus_species with 629 groups, obs with 1309 groups, phylo with 629 groups. The number of divergent transitions is 0. The maximum Rhat value is 1.003. |
| Longevity | The model was fitted using 1085 observations. It includes the following random effects: Genus_species with 471 groups, obs with 1085 groups, phylo with 471 groups. The number of divergent transitions is 0. The maximum Rhat value is 1.003. |
| Habitat | The model was fitted using 1439 observations. It includes the following random effects: Genus_species with 705 groups, obs with 1439 groups, phylo with 705 groups. The number of divergent transitions is 1. The maximum Rhat value is 1.005. |
| Sexual size dimorphism | The model was fitted using 848 observations. It includes the following random effects: Genus_species with 356 groups, obs with 848 groups, phylo with 356 groups. The number of divergent transitions is 0. The maximum Rhat value is 1.005. |
| Body mass | The model was fitted using 1438 observations. It includes the following random effects: Genus_species with 700 groups, obs with 1438 groups, phylo with 700 groups. The number of divergent transitions is 0. The maximum Rhat value is 1.002. |
| Breeding Season | The model was fitted using 1348 observations. It includes the following random effects: Genus_species with 675 groups, obs with 1348 groups, phylo with 675 groups. The number of divergent transitions is 1. The maximum Rhat value is 1.003. |
| Litter size | The model was fitted using 1438 observations. It includes the following random effects: Genus_species with 700 groups, obs with 1438 groups, phylo with 700 groups. The number of divergent transitions is 0. The |

**Table S5.** Summary of combinations of intraspecific variability in social organization, summarized by order.

| Taxa (order) | Common name | Species | Species included | Solitary | Mean IVSO |
| --- | --- | --- | --- | --- | --- |
| Afrosoricida | Tenrecs and golden moles | 56 | 3 | 67% | 0% |
| Artiodactyla | Even-toed ungulates | 335 | 117 | 22% | 22% |
| Carnivora | Carnivores | 297 | 101 | 48% | 10% |
| Chiroptera | Bats | 1,254 | 96 | 23% | 27% |
| Cingulata | Armadillos | 20 | 3 | 100% | 0% |
| Dermoptera | Colugos | 2 | 0 | -- | -- |
| Eulipotyphla | Hedgehogs, shrews, and moles | 494 | 16 | 45% | 28% |
| Hyracoidea | Hyraxes | 5 | 4 | 17% | 13% |
| Lagomorpha | Rabbits, hares, and pikas | 96 | 6 | 43% | 14% |
| Macroscelidea | Elephant shrews | 19 | 8 | 17% | 21% |
| Microbiotheriidae | Monito del monte | 1 | 0 | -- | -- |
| Perissodactyla | Odd-toed ungulates | 16 | 11 | 39% | 36% |
| Pholidota | Pangolins | 8 | 1 | 100% | -- |
| Pilosa | Anteaters and sloths | 10 | 6 | 100% | 3% |
| Primate | Primates | 445 | 215 | 3% | 19% |
| Proboscidea | Elephants | 2 | 2 | 24% | 13% |
| Rodentia | Rodents | 2,301 | 128 | 36% | 16% |
| Scandentia | Tree shrews | 23 | 5 | 18% | 13% |
| Sirenia | Sea cows | 5 | 3 | 25% | 4% |
| Tubulidentata | Aardvarks | 1 | 1 | 100% | 0% |

**Table S6.** Summary of combinations of social organization most commonly observed in species with IVSO. “PercentObservations” refers to the percent of study populations with IVSO that exhibited a given combination, while “PercentSpecies” refers to the perfect of species that exhibited a given combination, relative to all species in the dataset (i.e., placental mammals).

| Combinations of social organization | PercentObservations | PercentSpecies |
| --- | --- | --- |
| MFF + MMFF | 24.39 | 4.53 |
| MF + MFF | 9.76 | 1.89 |
| MF + Solitary | 6.71 | 1.39 |
| MMFF + Solitary | 6.10 | 1.13 |
| MF + MMFF | 6.10 | 1.13 |
| MMFF + SexSpecificGroup | 4.88 | 1.01 |
| MMF + MMFF | 4.27 | 0.88 |
| MFF + SexSpecificGroup | 3.05 | 0.50 |
| MF + MMFF + SexSpecificGroup + Solitary | 3.05 | 0.50 |
| SexSpecificGroup + Solitary | 3.05 | 0.63 |
| MF + MMF | 2.44 | 0.50 |
| MFF + Solitary | 2.44 | 0.50 |
| MF + SexSpecificGroup | 2.44 | 0.50 |
| MF + MMFF + Solitary | 1.83 | 0.38 |
| MFF + MMFF + Solitary | 1.83 | 0.25 |
| MF + MFF + Solitary | 1.83 | 0.38 |
| MMF + SexSpecificGroup | 1.83 | 0.38 |
| MFF + MMFF + SexSpecificGroup | 1.83 | 0.38 |
| MFF + MMFF + SexSpecificGroup + Solitary | 1.22 | 0.25 |
| MF + MFF + MMFF + Solitary | 1.22 | 0.25 |
| MFF + MMF | 1.22 | 0.25 |
| MF + MMF + MMFF | 1.22 | 0.25 |
| MF + MFF + MMF | 1.22 | 0.25 |
| MMF + SexSpecificGroup + Solitary | 0.61 | 0.13 |
| MFF + SexSpecificGroup + Solitary | 0.61 | 0.13 |
| MMFF + SexSpecificSolitaryM | 0.61 | 0.13 |
| MF + MFF + MMFF + SexSpecificGroup | 0.61 | 0.13 |
| MF + MFF + MMFF | 0.61 | 0.13 |
| MFF + MMF + MMFF + SexSpecificGroup + Solitary | 0.61 | 0.13 |
| MF + MFF + MMFF + SexSpecificGroup + Solitary | 0.61 | 0.13 |
| MMFF + SexSpecificGroup + Solitary | 0.61 | 0.13 |
| MF + MFF + SexSpecificGroup + Solitary | 0.61 | 0.13 |
| MF + MFF + MMF + MMFF + SexSpecificGroup | 0.61 | 0.13 |

## References

1 Clutton-Brock, T. Social evolution in mammals. Science 373, eabc9699, doi:10.1126/science.abc9699 (2021).

2 Lukas, D. & Clutton-Brock, T. H. The evolution of social monogamy in mammals. Science 341, 526–530, doi:10.1126/science.1238677 (2013).

3 Koenig, A., Scarry, C. J., Wheeler, B. C. & Borries, C. Variation in grouping patterns, mating systems and social structure: what socio-ecological models attempt to explain. Philosophical Transactions of the Royal Society B: Biological Sciences 368, doi:10.1098/rstb.2012.0348 (2013).

4 Shultz, S., Opie, C. & Atkinson, Q. D. Stepwise evolution of stable sociality in primates. Nature 479, 219–222, doi:http://www.nature.com/nature/journal/v479/n7372/abs/nature10601.html#supplementary-information (2011).

5 Kappeler, P. M. & Pozzi, L. Evolutionary transitions toward pair living in nonhuman primates as stepping stones toward more complex societies. Science Advances 5, eaay1276, doi:10.1126/sciadv.aay1276 (2019).

6 Olivier, C.-A., Jaeggi, A. V., Hayes, L. D. & Schradin, C. Revisiting the components of Macroscelidea social systems: Evidence for variable social organization, including pair-living, but not for a monogamous mating system. Ethology 128, 383–394, 10.1111/eth.13271 (2022).

7 Jaeggi, A. V., Miles, M. I., Festa-Bianchet, M., Schradin, C. & Hayes, L. D. Variable social organization is ubiquitous in Artiodactyla and probably evolved from pair-living ancestors. Proceedings of the Royal Society B: Biological Sciences 287, 20200035, doi:10.1098/rspb.2020.0035 (2020).

8 Olivier, C.-A. et al. Primate social organization evolved from a flexible pair-living ancestor. Proceedings of the National Academy of Sciences 121, e2215401120, doi:10.1073/pnas.2215401120 (2024).

9 Qiu, J., Olivier, C. A., Jaeggi, A. V. & Schradin, C. The evolution of marsupial social organization. Proceedings of the Royal Society B: Biological Sciences 289, 20221589, doi:10.1098/rspb.2022.1589 (2022).

10 Schradin, C. Comparative studies need to rely both on sound natural history data and on excellent statistical analysis. Royal Society Open Science 4: 170346, 10.1098/rsos.170346 (2017).

11 Brusatte, S. The Rise and Reign of the Mammals. (Picador, 2022).

12 Weaver, L. N. et al. Early mammalian social behaviour revealed by multituberculates from a dinosaur nesting site. Nature Ecology & Evolution, doi:10.1038/s41559-020-01325-8 (2020).

13 Ladeveze, S., de Muizon, C., Beck, R. M. D., Germain, D. & Cespedes-Paz, R. Earliest evidence of mammalian social behaviour in the basal Tertiary of Bolivia. Nature 474, 83–86, doi:http://www.nature.com/nature/journal/v474/n7349/abs/10.1038-nature09987-unlocked.html#supplementary-information (2011).

14 Jasinoski, S. C. & Abdala, F. Aggregations and parental care in the Early Triassic basal cynodonts Galesaurus planiceps and Thrinaxodon liorhinus. PeerJ 5, e2875, doi:10.7717/peerj.2875 (2017).

15 Kappeler, P. M. A framework for studying social complexity. Behavioral Ecology and Sociobiology 73, 13, doi:10.1007/s00265-018-2601-8 (2019).

16 Fernandez-Duque, E., Huck, M., Van Belle, S. & Di Fiore, A. The evolution of pair-living, sexual monogamy, and cooperative infant care: Insights from research on wild owl monkeys, titis, sakis, and tamarins. American Journal of Physical Anthropology 171, 118–173, 10.1002/ajpa.24017 (2020).

17 Bales, K. L. et al. What is a pair bond? Hormones and Behavior 136, 105062, 10.1016/j.yhbeh.2021.105062 (2021).

18 Huck, M., Di Fiore, A. & Fernandez-Duque, E. Of apples and oranges? The evolution of “monogamy” in non-human primates. Frontiers in Ecology and Evolution 7, doi:10.3389/fevo.2019.00472 (2020).

19 Kvarnemo, C. Why do some animals mate with one partner rather than many? A review of causes and consequences of monogamy. Biological Reviews 93, 1795–1812, doi:10.1111/brv.12421 (2018).

20 Schradin, C., Hayes, L. D., Pillay, N. & Bertelsmeier, C. The evolution of intraspecific variation in social organization. Ethology 124, 527–536, doi:10.1111/eth.12752 (2018).

21 Rubenstein, D. R. & Abbot, P. in Comparative Social Evolution (eds Dustin R. Rubenstein & Patrick Abbot) 1–18 (Cambridge University Press, 2017).

22 Crook, J. H. & Gartlan, J. S. Evolution of primate societies. Nature 210, 1200–1203 (1966).

23 Emlen, S. T. & Oring, L. W. Ecology, secual selection, and the evolution of mating systems. Science 197, 215–223 (1977).

24 Terborgh, J. & Janson, C. H. The socioecology of primate groups. Annual Review of Ecology and Systematics 17, 111–136, doi:10.1146/annurev.es.17.110186.000551 (1986).

25 Krause, J. & Ruxton, G. D. Living in Groups. (Oxford University Press, 2002).

26 Markham, A. C. & Gesquiere, L. R. Costs and benefits of group living in primates: an energetic perspective. Philosophical Transactions of the Royal Society B: Biological Sciences 372, doi:10.1098/rstb.2016.0239 (2017).

27 Van Schaik, C. P. & Hooff, J. A. R. A. M. V. On the Ultimate Causes of Primate Social Systems. Behaviour 85, 91–117 (1983).

28 Jordan, N. R. et al. Priority of access to food and its influence on social dynamics of an endangered carnivore. Behavioral Ecology and Sociobiology 76, 13, doi:10.1007/s00265-021-03115-z (2022).

29 Wrangham, R. W. An ecological model of female-bonded primate groups. Behaviour 75, 262–300 (1980).

30 White, F. J. Seasonality and Socioecology: The Importance of Variation in Fruit Abundance to Bonobo Sociality. International Journal of Primatology 19, 1013–1027, doi:10.1023/A:1020374220004 (1998).

31 Thierry, B. External and internal constraints in socioecology. Phil Trans R Soc B (2013).

32 Gittleman, J. L. & Valkenburgh, B. V. Sexual dimorphism in the canines and skulls of carnivores: effects of size, phylogency, and behavioural ecology. Journal of Zoology 242, 97–117, 10.1111/j.1469-7998.1997.tb02932.x (1997).

33 Tombak, K. J., Hex, S. B. S. W. & Rubenstein, D. I. New estimates indicate that males are not larger than females in most mammal species. Nature Communications 15, 1872, doi:10.1038/s41467-024-45739-5 (2024).

34 Jeschke, J., Gabriel, W. & Kokko, H. in Encyclopedia of Ecology 3113-3122 (Elsevier, 2008).

35 Silk, J. B. The adaptive value of sociality in mammalian groups. Philos Trans R Soc Lond B Biol Sci 362, 539–559, doi:10.1098/rstb.2006.1994 (2007).

36 Valomy, M., Hayes, L. D. & Schradin, C. Social organization in Eulipotyphla: evidence for a social shrew. Biology Letters 11, doi:10.1098/rsbl.2015.0825 (2015).

37 Luo, Z.-X., Yuan, C.-X., Meng, Q.-J. & Ji, Q. A Jurassic eutherian mammal and divergence of marsupials and placentals. Nature 476, 442–445, doi:10.1038/nature10291 (2011).

38 Warren, W. C. et al. Genome analysis of the platypus reveals unique signatures of evolution. Nature 453, 175–183, doi:10.1038/nature06936 (2008).

39 Brink, A. S. Speculations on some advanced mammalian characteristics in the higher mammal-like reptiles. Palaeontologia Africana 4, 77–96 (1956).

40 Groenewald, G. H., Welman, J. & Maceachern, J. Vertebrate burrow complexes from the early Triassic Cynognathus zone (Driekoppen formation, Beaufort group) of the Karoo basin, South Africa. PALAIOS 16, 148–160, doi:10.2307/3515526 (2001).

41 Griffith, S. C., Owens, I. P. F. & Thuman, K. A. Extra pair paternity in birds: a review of interspecific variation and adaptive function. Mol. Ecol. 11, 2195–2212 (2002).

42 Kempenaers, B. Mating systems in birds. Current Biology 32, R1115–R1121, 10.1016/j.cub.2022.06.066 (2022).

43 Flannery, T. F., Rich, T. H., Vickers-Rich, P., Veatch, E. G. & Helgen, K. M. The Gondwanan Origin of Tribosphenida (Mammalia). Alcheringa: An Australasian Journal of Palaeontology 46, 277–290, doi:10.1080/03115518.2022.2132288 (2022).

44 Novacek, M. J. et al. Epipubic bones in eutherian mammals from the Late Cretaceous of Mongolia. Nature 389, 483–486, doi:10.1038/39020 (1997).

45 Sandell, M. I. Female aggression and the maintenance of monogamy: Female behaviour predicts male mating status in European starlings. Proc R Soc Lond B 265, 1307–1311 (1998).

46 Stockley, P. & Bro-Jørgensen, J. Female competition and its evolutionary consequences in mammals. Biological Reviews 86, 341–366, 10.1111/j.1469-185X.2010.00149.x (2011).

47 Tucker, M. A., Ord, T. J. & Rogers, T. L. Evolutionary predictors of mammalian home range size: body mass, diet and the environment. Global Ecology and Biogeography 23, 1105–1114, 10.1111/geb.12194 (2014).

48 Schradin, C. & Ancel, A. in Encyclopedia of Animal Cognition and Behavior (eds Jennifer Vonk & Todd Shackelford) 1–5 (Springer International Publishing, 2019).

49 Funston, G. F. et al. The origin of placental mammal life histories. Nature, doi:10.1038/s41586-022-05150-w (2022).

50 Pollard, K. A. & Blumstein, D. T. Social group size predicts the evolution of individuality. Current Biology 21, 413–417 (2011).

51 Mocha, Y. B. et al. Advancing cooperative breeding research with a peer-reviewed and “live” Cooperative-Breeding Database (Co-BreeD). bioRxiv, 2024.2004.2026.591342, doi:10.1101/2024.04.26.591342 (2024).

52 Taborsky, M. Sneakers, satellites, and helpers: parasitic and cooperative behavior in fish reproduction. Adv Study Behav 23, 1–100 (1994).

53 Emlen, S. T. An evolutionary theory of the family. Proc Natl Acad Sci USA 92, 8092–8099 (1995).

54 Myers, T. S. & Fiorillo, A. R. Evidence for gregarious behavior and age segregation in sauropod dinosaurs. Palaeogeography, Palaeoclimatology, Palaeoecology 274, 96–104, 10.1016/j.palaeo.2009.01.002 (2009).

55 Ostrom, J. H. Were some dinosaurs gregarious? Palaeogeography, Palaeoclimatology, Palaeoecology 11, 287–301, 10.1016/0031-0182(72)90049-1 (1972).

56 Botfalvai, G., Prondvai, E. & Ősi, A. Living alone or moving in herds? A holistic approach highlights complexity in the social lifestyle of Cretaceous ankylosaurs. Cretaceous Research 118, 104633, 10.1016/j.cretres.2020.104633 (2021).

57 Makuya, L., Pillay, N., Sangweni, S. P. & Schradin, C. Tolerant mothers: aggression does not explain solitary living in the bush Karoo rat. Proceedings of the Royal Society B: Biological Sciences 291, 20241534, doi:10.1098/rspb.2024.1534 (2024).

58 Makuya, L. & Schradin, C. Costs and benefits of solitary living in mammals. Journal of Zoology 323, 9–18, 10.1111/jzo.13145 (2024).

59 Upham, N. S., Esselstyn, J. A. & Jetz, W. Inferring the mammal tree: Species-level sets of phylogenies for questions in ecology, evolution, and conservation. PLOS Biology 17, e3000494, doi:10.1371/journal.pbio.3000494 (2019).

60 Makuya, L., Olivier, C.-A. & Schradin, C. Field studies need to report essential information on social organisation – independent of the study focus. Ethology 128, 268–274, 10.1111/eth.13249 (2022).

61 Grueter, C. C. et al. Multilevel organisation of animal sociality. Trends in Ecology & Evolution 35, 834–847, 10.1016/j.tree.2020.05.003 (2020).

62 Goldizen, A. W. A comparative perspective on the evolution of tamarin and marmoset social systems. Int J Primatol 11, 63–83 (1990).

63 Walmsley, S. F. et al. (data.InDoRES, 2025).

64 Schradin, C. Intraspecific variation in social organization by genetic variation, developmental plasticity, social flexibility or entirely extrinsic factors. Philosophical Transactions of the Royal Society B-Biological Sciences 368, doi:10.1098/rstb.2012.0346, doi:10.1098/rstb.2012.0346 (2013).

65 Dalerum, F. Phylogenetic reconstruction of carnivore social organizations. Journal of Zoology 273, 90–97, doi:10.1111/j.1469-7998.2007.00303.x (2007).

66 Wilson, D. E. & Mittermeier, R. A. Handbook of the Mammals of the World. (2018).

67 Jones, K. E. et al. PanTHERIA: a species-level database of life history, ecology, and geography of extant and recently extinct mammals. Ecology 90, 2648–2648, 10.1890/08-1494.1 (2009).

68 De Magelhaes, J. P. & Costa, J. A database of vertebrate longevity records and their relation to other life-history traits. Journal of Evolutionary Biology 22, 1770–1774, 10.1111/j.1420-9101.2009.01783.x (2009).

69 Hadfield, J. D. & Nakagawa, S. General quantitative genetic methods for comparative biology: phylogenies, taxonomies and multi-trait models for continuous and categorical characters. Journal of Evolutionary Biology 23, 494–508, doi:10.1111/j.1420-9101.2009.01915.x (2010).

70 McElreath, R. Statistical rethinking: A Bayesian course with examples in R and Stan (2nd editio). (CRC Press, 2020).

71 Lemoine, N. P. Moving beyond noninformative priors: why and how to choose weakly informative priors in Bayesian analyses. Oikos 128, 912–928, 10.1111/oik.05985 (2019).

72 Garland, T., Jr. & Ives, A. R. Using the Past to Predict the Present: Confidence Intervals for Regression Equations in Phylogenetic Comparative Methods. Am Nat 155, 346–364, doi:10.1086/303327 (2000).

73 Álvarez-Carretero, S. et al. A species-level timeline of mammal evolution integrating phylogenomic data. Nature 602, 263–267, doi:10.1038/s41586-021-04341-1 (2022).

74 Carlisle, E., Janis, C. M., Pisani, D., Donoghue, P. C. J. & Silvestro, D. A timescale for placental mammal diversification based on Bayesian modeling of the fossil record. Current Biology, doi:10.1016/j.cub.2023.06.016 (2023).

75 Foley, N. M. et al. A genomic timescale for placental mammal evolution. Science 380, eabl8189, doi:10.1126/science.abl8189 (2023).

76 Ji, Q. et al. The earliest known eutherian mammal. Nature 416, 816–822, doi:10.1038/416816a (2002).

77 R: A Language and Environment for Statistical Computing. (Vienna, Austria, 2022).

78 Bürkner, P.-C. brms: An R Package for Bayesian Multilevel Models Using Stan. 2017 80, 28, doi:10.18637/jss.v080.i01 (2017).

79 Landau, W. M. The stantargets R package: a workflow framework for efficient reproducible Stan-powered Bayesian data analysis pipelines. Journal of Open Source Software 6, 3139, 10.21105/joss.03193 (2021).

## References

1 10.5281/zenodo.15007505, M. D. D. M. D. D. v. D. s. Z. (2025).

2 Makuya, L., Olivier, C.-A. & Schradin, C. Field studies need to report essential information on social organisation – independent of the study focus. Ethology 128, 268–274, 10.1111/eth.13249 (2022).

3 Jones, K. E. et al. PanTHERIA: a species-level database of life history, ecology, and geography of extant and recently extinct mammals. Ecology 90, 2648–2648, 10.1890/08-1494.1 (2009).

4 Wilson, D. E. & Mittermeier, R. A. Handbook of the Mammals of the World. (2018).

5 De Magelhaes, J. P. & Costa, J. A database of vertebrate longevity records and their relation to other life-history traits. Journal of Evolutionary Biology 22, 1770–1774, 10.1111/j.1420-9101.2009.01783.x (2009).

